# Effective mechanical potential of cell–cell interaction in tissues harboring cavity and in cell sheet toward morphogenesis

**DOI:** 10.1101/2023.08.28.555208

**Authors:** Hiroshi Koyama, Hisashi Okumura, Tetsuhisa Otani, Atsushi M. Ito, Kazuyuki Nakamura, Kagayaki Kato, Toshihiko Fujimori

## Abstract

Measuring mechanical forces of cell–cell interactions is important for studying morphogenesis in multicellular organisms. We previously reported an image-based statistical method for inferring effective mechanical potentials of pairwise cell–cell interactions by fitting cell tracking data with a theoretical model. However, whether this method is applicable to tissues with non-cellular components such as cavities remains elusive. Here we evaluated the applicability of the method to cavity-harboring tissues. Using synthetic data generated by simulations, we found that the effect of expanding cavities was added to the pregiven potentials used in the simulations, resulting in the inferred effective potentials having an additional repulsive component derived from the expanding cavities. Interestingly, simulations by using the effective potentials reproduced the cavity-harboring structures. Then, we applied our method to the mouse blastocysts, and found that the inferred effective potentials can reproduce the cavity-harboring structures. Pairwise potentials with additional repulsive components were also detected in two-dimensional cell sheets, by which curved sheets including tubes and cups were simulated. We conclude that our inference method is applicable to tissues harboring cavities and cell sheets, and the resultant effective potentials are useful to simulate the morphologies.

## Introduction

In multicellular living systems, various three-dimensional morphologies are emerged, which is thought to be primarily dependent on the mechanical properties of the constituent cells (Steinberg, 2007; Maître et al., 2012, 2015; Guillot and Lecuit, 2013; Fletcher et al., 2014; Bazellières et al., 2015; Petridou and Heisenberg, 2019). Measurement of cellular mechanical parameters is inevitable for understanding morphogenetic events. On the other hand, morphogenetic events are also influenced by non-cellular structures such as cavities, extracellular matrix, etc (Style et al., 2014; Wang et al., 2018). Therefore, it is challenging to identify all components contributing to morphogenetic events, and then, to measure their mechanical forces especially under three-dimensional situations.

Image-based methods for inferring mechanical forces in tissues have been developed, where mathematical models are assumed which are fitted to microscopic images (Miyoshi et al., 2006; Brodland et al., 2010; Ishihara and Sugimura, 2012; Koyama et al., 2012, 2023; Merkel and Manning, 2017; Kondo et al., 2018). These methods include inference of cell–cell junction tensions in two-dimensional epithelia, cell surface tensions or bending rigidities, mechanical potential of pairwise cell–cell interactions, etc. Basically, these methods do not assume non-cellular structures in their mathematical models, and, it is unknown whether these methods are applicable to tissues with non-cellular components.

We previously reported a method for inferring mechanical potential of pairwise cell–cell interactions (Koyama et al., 2023). The pairwise potentials are composed of attractive and repulsive forces of cell–cell interactions. The attractive forces are at least partially derived from cell–cell adhesion forces mediated by cell adhesion molecules (Fig. 1A-B), whereas the repulsive forces are from excluded volume effect of cells (Drasdo et al., 1995; Forgacs and Newman, 2005; Steinberg, 2007; Maître et al., 2012; Guillot and Lecuit, 2013; Yong et al., 2015; Merkel and Manning, 2017). The summation of these forces would be dependent on cell–cell distance (Fig. 1C), by which the pairwise potential energy is calculated. We previously showed that, by applying our method to the mouse and *C. elegans* early embryos which did not still harbor clear cavities or thick extracellular matrix, we obtained mechanical potential of pairwise interactions (Koyama et al., 2023). The profiles of the pairwise potentials were quantitatively different among the tissues, by which the morphologic features of the corresponding tissues were reproduced.

**Figure 1:**
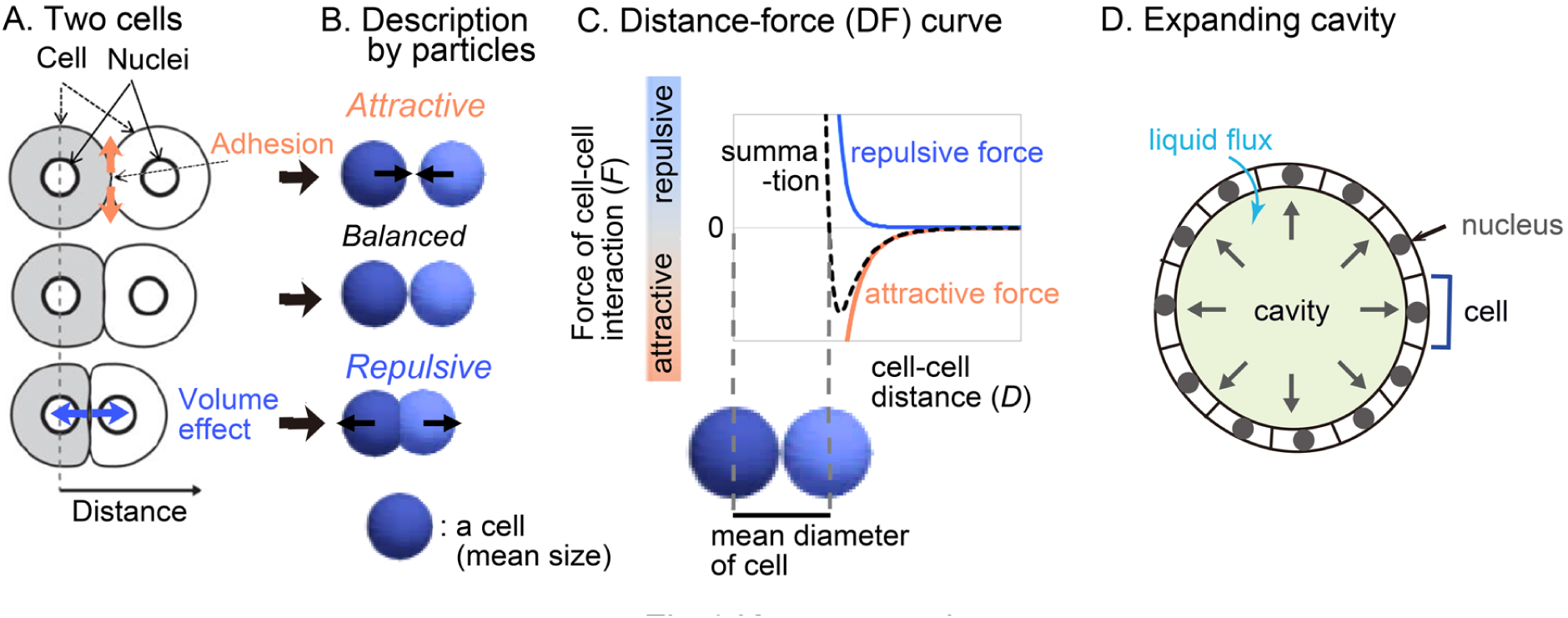
Overview of strategy for inferring the effective potential of cell–cell interactions A. Microscopic forces contributing to cell–cell interactions are exemplified. B. The microscopic forces are approximated as attractive/repulsive forces where cells are described as particles with the mean size. C. Both the repulsive and attractive forces are provided as cell–cell distance-dependent functions, and the summation (black broken line) is the distance–force curve of the two particles. The relationship between the curve and the mean diameters of the cells is shown. D. A cavity-harboring tissue is illustrated with liquid flux into the cavity.

In the present study, we evaluated whether our inference method is applicable to cavity-harboring tissues such as the mouse blastocysts (Fig. 1D). First, we theoretically analyzed the effect of liquid cavities by using synthetic data which were generated by simulations under pregiven cell–cell forces and cavities. In the case where the cavities are expanding, the inferred effective pairwise forces of cell–cell interactions became significantly different from the pregiven forces, where an additional repulsive component was detected. But we showed that the repulsive component was derived from the expanding cavities, and that the inferred effective pairwise forces were the summation of both the pregiven forces and the effect of the expanding cavities. Therefore, simulations using inferred pairwise forces successfully reproduced the cavity-harboring structures. Application of our method to the mouse blastocysts also resulted in the same conclusion. Because the effect of cavities on the inferred effective forces is additive, we may evaluate differentials of cell–cell interaction forces between two tissues such as wild-type and mutant ones, if one can assume that the effect of the cavities is equivalent between the two tissues. Finally, we systematically analyzed what kind of profiles of the effective pairwise potentials are critical for forming cavity-harboring structures, which were presented as phase diagrams, and expanded the analyses to curved sheets.

## Theory; force inference procedure with simulation model

### 2.1 Overview

In our inference method, a cell-particle model is assumed where attractive/repulsive forces of cell–cell interaction is considered (Fig. 1B-C), and the mathematical model is fitted to nuclear tracking data (i.e., *xyz*-coordinates of each nucleus with the time series) as described previously (Koyama et al., 2023). Both the modeled particles and the nuclei were assumed as point particles. Briefly, the fitting procedure is based on a minimization of the differences between the *xyz*-coordinates of point particles in the mathematical model and in the nuclear tracking data, with an additional cost function for cell–cell distance-dependent weight. The fitting parameters are the force values for each pair of two interacting cells for each time frame, and then, by binned averaging the inferred forces along cell–cell distances (*D*), a distance–force curve (*F*_C–C_(*D*)) is calculated. In the model to be fitted, we do not explicitly consider non-cellular structures exerting forces on cells. It can be argued that, if tissues harbor non-cellular structures, the model should assume the non-cellular structures. But, due to high degrees of freedom in such models, the fitting problems would become difficult to be solved in general. We intended to analyze whether valuable information can be obtained when our inference method is applied to tissues with cavities (Fig. 1D). To this aim, we first generated simulation data considering both cell–cell interaction forces and cavities, to which we applied our inference method.

### 2.2 Simulation model

In our simulations based on particulate cells, the cell–cell interaction forces were set to be provided by the Lennard-Jones (LJ) potential (i.e., the pregiven forces) in a similar manner to our previous work (Koyama et al., 2023). Briefly, due to the low Reynolds number in the case of cellular-level phenomena, the inertial force is negligible. Viscous drag force or frictional forces are provided from surrounding liquid medium or tissues (Forgacs and Newman, 2005; Basan et al., 2011; Fletcher et al., 2014; Li et al., 2017). If the frictional forces are provided from non-moving surrounding medium or substrates, the equation of a particle motion may be written as *V*_C*|p*_ = *F*_C*|p*_ /*γ* for each particle, where *V*_C*|p*_ is the velocity of the *p*th particle, *F*_C*|p*_ is the summation of cell–cell interaction forces exerted on the *p*th particle, and *γ* is the coefficient of viscous drag and frictional forces provided from surrounding medium or substrates; we call it “Model-A”. This assumption has been used in both particle-based models and vertex models (a standard model for epithelial cells) including simulations for the blastocysts (Honda et al., 2008a; Krupinski et al., 2011; Fletcher et al., 2014; Li et al., 2017; Nissen et al., 2017, 2018). On the other hand, if viscous frictional forces are dominantly provided from neighboring particles in motion, relative velocities between particles become important, and the equation of a particle motion may be written as follows; 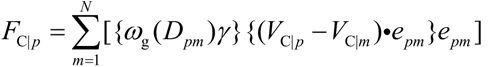, where *m* is the ID of particles interacting with the *p*th particle, *V*_C|*m*_ is the velocity of the *m*th particle, and *N* is the total number of the interacting particles. *D_pm_* is the distance between the *p*th and *m*th particles, *ω*_g_(*D_pm_*) is a distance-dependent weight for *γ*, and *e_pm_* is the unit vector from the xyz-coordinates of *m*th to *p*th particles. *ω*_g_(*D_pm_*) decays along *D_pm_*. We call it “Model-B”. This assumption has been used in particle-based models especially for three-dimensionally packed cell aggregates (Nikunen et al., 2003; Basan et al., 2011; Podewitz et al., 2015). A similar assumption based on relative velocities was also adopted in a three-dimensional vertex model (Okuda et al., 2015b). Note that, in our inference method, equivalent force values were obtained between the above two assumptions (Koyama et al., 2023).

### 2.3 Force inference method

In our force inference method, the above model is fitted to nuclear tracking data. The cost function to be minimized is as follows; *G* = *G_xyz_* + *G_F_0__*, where *G_xyz_* is the difference between *xyz*-coordinates of each particle of *in vivo* data and particle simulations, and *G_F_0__* is the additional cost function for cell–cell distance-dependent weight as previously defined (Koyama et al., 2023). The definition is described in Supplementary information. Our inference method was validated by applying this method to simulation data generated under pregiven forces (Koyama et al., 2023).

## Results

### 3.1 Inference of effective pairwise potential in synthetic data with cavity

To generate simulation data with expanding cavities, we modeled them as follows. In real tissues, cavities are generated by secreting liquid from cells. The cell layers surrounding the cavities are sealed by the tight junctions, and thus the pressure of the cavities push the cells outward (Fig. 1D) (Moriwaki et al., 2007; Wang et al., 2018; Dumortier et al., 2019). In simulations, we implemented expanding spherical cavities as temporally-evolved spatial constraints; the particulate cells cannot penetrate into the cavities (Fig. S1A-B and movieS1). Although this constraint was not functional to prevent the particles from going outside of the cavities (Fig. S1B), the particles were almost constrained to move on the surfaces of the cavities.

Fig. 2 is the simulation results based on Model-A, and Fig. 2A-i shows the snapshot of the simulations where the spherical cavity was expanding from the radius = 9.6 to 12.6μm. Fig. 2A-ii shows inferred effective distance–force (DF) curves under three different conditions of the expanding cavities; the inferred DF curves were not consistent with the DF curves from the LJ potential (Fig. 2A-ii, compare with the black broken line), where repulsive components were strongly detected especially at distant regions, which we call distant energy barrier (DEB). The strength of the repulsive components seemed to be positively related to the expansion rates. Under the higher expansion rate (Fig. 2A; 9.7->14.2), attractive components were diminished in the DF curves, meaning that the attractive forces derived from the LJ potential were effectively cancelled. In contrast, under the conditions of non-expanding cavities, the inferred DF curves were consistent with the LJ potential (Fig. S1C), probably because the non-expanding cavities did barely push the particles outwardly. Similar repulsive components were detected in systems with both larger cavities and larger numbers of particles (Fig. S1D; particle number = 150, radius of cavity = ∼ 50μm). Moreover, our results in Fig. 2A were scalable; i.e., if the sizes of the cavities, and both the distance and the force in the LJ potential were set to be multiplied by the same value in the simulations (e.g., ×5), inferred DF curves were also multiplied by the same value (Fig. S1E; e.g., radius of cavity = ∼40μm).

**Figure 2:**
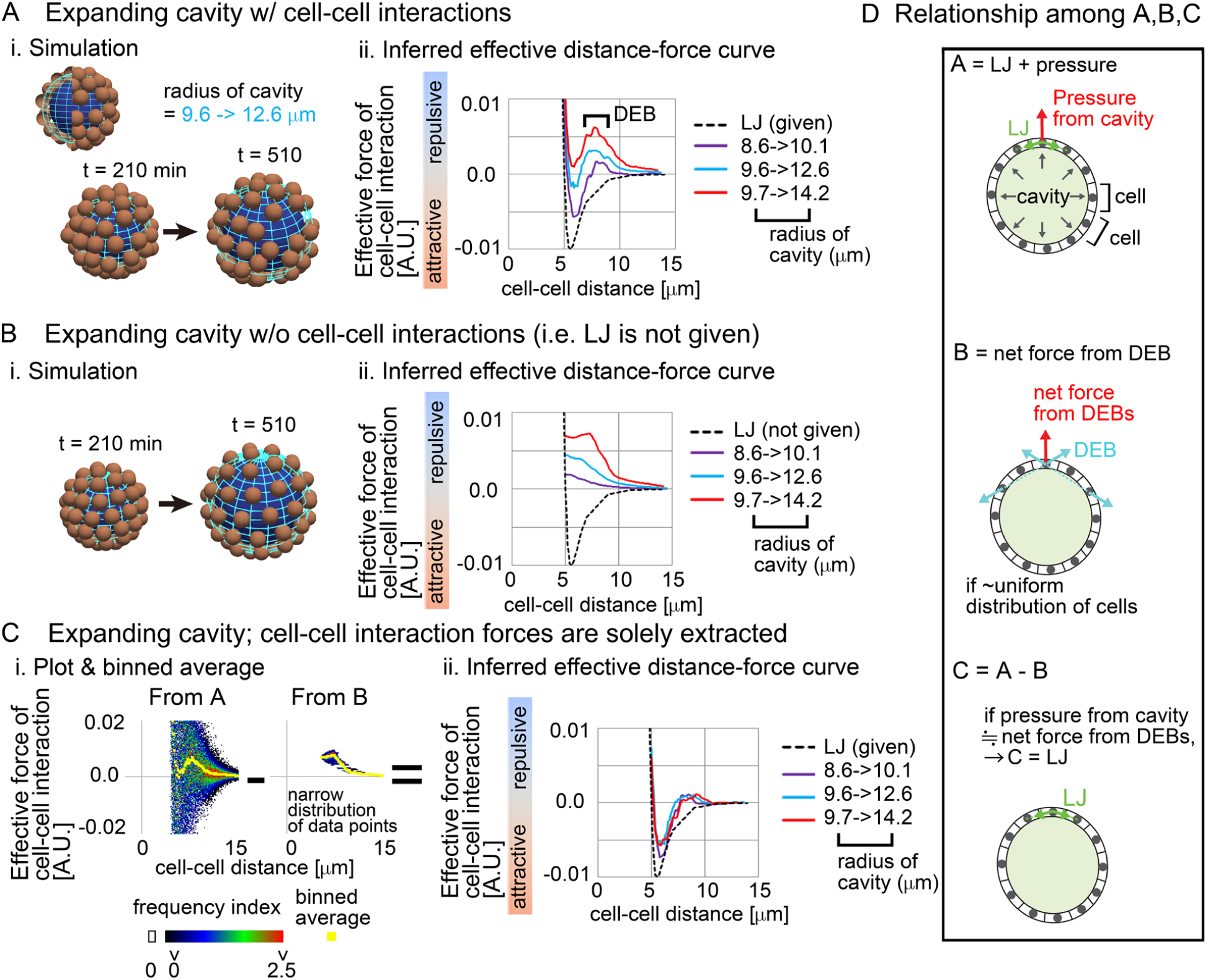
Influence of external constraints to effective forces of cell–cell interaction Simulations were performed on the basis of a distance–force (DF) curve obtained from the Lennard-Jones potential (LJ). A. Expanding cavities are assumed in the particle model. i) An example of simulations with the change in the radius of the cavity. The particles are of the mean size of cells. ii) Inferred effective DF curves with the distant energy barriers (DEBs). The pregiven DF curve (LJ) is shown in a broken black line. The numbers of the particles (= 64) are the same among the different cavities. To exclude the possible influence of initial particle configurations on the force inference, particle tracking data from *t* = 210 min but not 0 min was applied to the force inference. B. Expanding cavities are assumed whereas cell–cell interaction forces are not introduced. i) An example of simulations. ii) Inferred effective DF curves. C. Reconstruction of cell–cell interaction forces under expanding cavities. i) All the data points in A and B under the radius of the cavity = 9.7 -> 14.2μm are plotted on distance– force graph as a heat map, and the binned averages were calculated (yellow) which correspond to the DF curves in A and B. Each heat map is derived from forces inferred from one simulation. ii) DF curves obtained by subtracting the DF curves in B from those in A: i.e., A – B as exemplified in C-i. The definition of the heat map is described in Supplementary information. D. The subtraction performed in C is schematically explained in the relationship among A, B, and C. See the main text. Related figure: Figure S1 (simulation procedures and inferred DF curves under both non-expanding cavities and larger expanding cavities with larger number of particles). Related movie: Movie S1 (simulation)

We then examined whether this modification is meaningless or is a result of a correct reflection of the expanding cavities. By performing simulations under conditions with the expanding cavities but without the LJ potential (i.e., without both the attractive and repulsive forces of cell–cell interaction) (Fig. 2B-i), we solely extracted the influence of the expanding cavities. Fig. 2B-ii shows that the inferred effective DF curves were exclusively composed of repulsive components; i.e., attractive components did not appear. Then, we checked the raw data where all data points were plotted on the distance–force space; in Fig. 2C-i, “From B”, a heat map of the data points is shown. Force values calculated by binned averaging are also shown in yellow, which correspond to the DF curve in Fig. 2B-ii (red line). Interestingly, all the data points were closely located around the DF curve (“From B”), implying that the DF curve is a good approximation of the expanding cavities. Probably, due to the effect of the expanding cavities which causes the increases in distances among the particles, pairwise forces of particle–particle interactions became apparently repulsive.

Finally, by using the above information, we examined whether the correct DF curve (i.e., LJ potential) can be reconstructed. We subtracted the DF curves in Fig. 2B from those in Fig. 2A (Fig. 2C-i, “From A – From B”), and the resultant DF curves were well consistent with the LJ potential under any expansion rates of the cavities (Fig. 2C-ii). These results suggest that the effects of the expanding cavities can be additive to effective DF curves.

In addition, the maximum values of the attractive forces in Fig. 2C-ii were 50∼70% of that in the LJ potential. According to our previous study (Koyama et al., 2023), equivalent decreases in the inferred attractive forces were seen in systems without cavities, and sampling intervals also affected the values of attractive forces. Therefore, we think that the decreases are derived from a measurement error. The shorter the sampling interval, the more accurate the inferred forces (Koyama et al., 2023). However, in real three-dimensional tissues, we are usually unable to set very short sampling intervals due to technical difficulties, so we typically set it to be ≥3 min in the case of the mouse embryos as shown later. In Fig. 2, we adopted a sampling interval equivalent to the case of the mouse embryos.

We interpret the above results as described in Fig. 2D. A cell receives forces from cell–cell interactions (“LJ” in Fig. 2D) and from the pressure of the cavity, where the vector of the pressure is perpendicular to the surface of the cavity (Fig. 2D, top panel). The repulsive forces detected as cell–cell interactions in Fig. 2A are described in Fig. 2D, middle panel (light blue vectors), and the vector of the net force (a red vector) for a cell is perpendicular to the surface of the cavity, if the cells are almost uniformly distributed. Therefore, the force provided by the pressure of the cavity can be equivalent to the net force from the DEBs (Fig. 2D, bottom panel), leading to the successful reconstruction of the LJ potential.

We conclude that the expanding cavities can be approximated as effective pairwise potentials. In general, effective pairwise forces/potentials are the summation of different components as explained in Fig. 1C. External factors are one of the components to be summed up in molecular and colloidal sciences: e.g., by considering hydration of particles by solvents, effective pairwise potentials are modified so as to contain energy barriers around distant regions (Pettitt and Rossky, 1986; Israelachvili, 2011), and such effective potentials have been used to simulate behaviors of the systems. We will show the simulation outcome from the above inferred effective DF curve later.

### 3.2 Effective forces in mouse blastocyst contained DEB

We investigated whether a repulsive component appeared in effective forces inferred from the mouse blastocyst (Fig. 3A, illustration). An inner cavity is formed through liquid secretion from the cells, and the embryos expand while maintaining their spherical shape (Moriwaki et al., 2007; Wang et al., 2018; Dumortier et al., 2019) (Fig. 3A, bright field and cross-section). Trophectoderm (TE) cells form a single-cell-layered structure at the outermost surface of the embryo, whereas the cells of the inner cell mass (ICM) were tightly assembled, forming a cluster (Fig. 3A, illustration). We performed cell tracking of both TE and ICM (Fig. 3A, tracking) and inferred effective forces.

**Figure 3:**
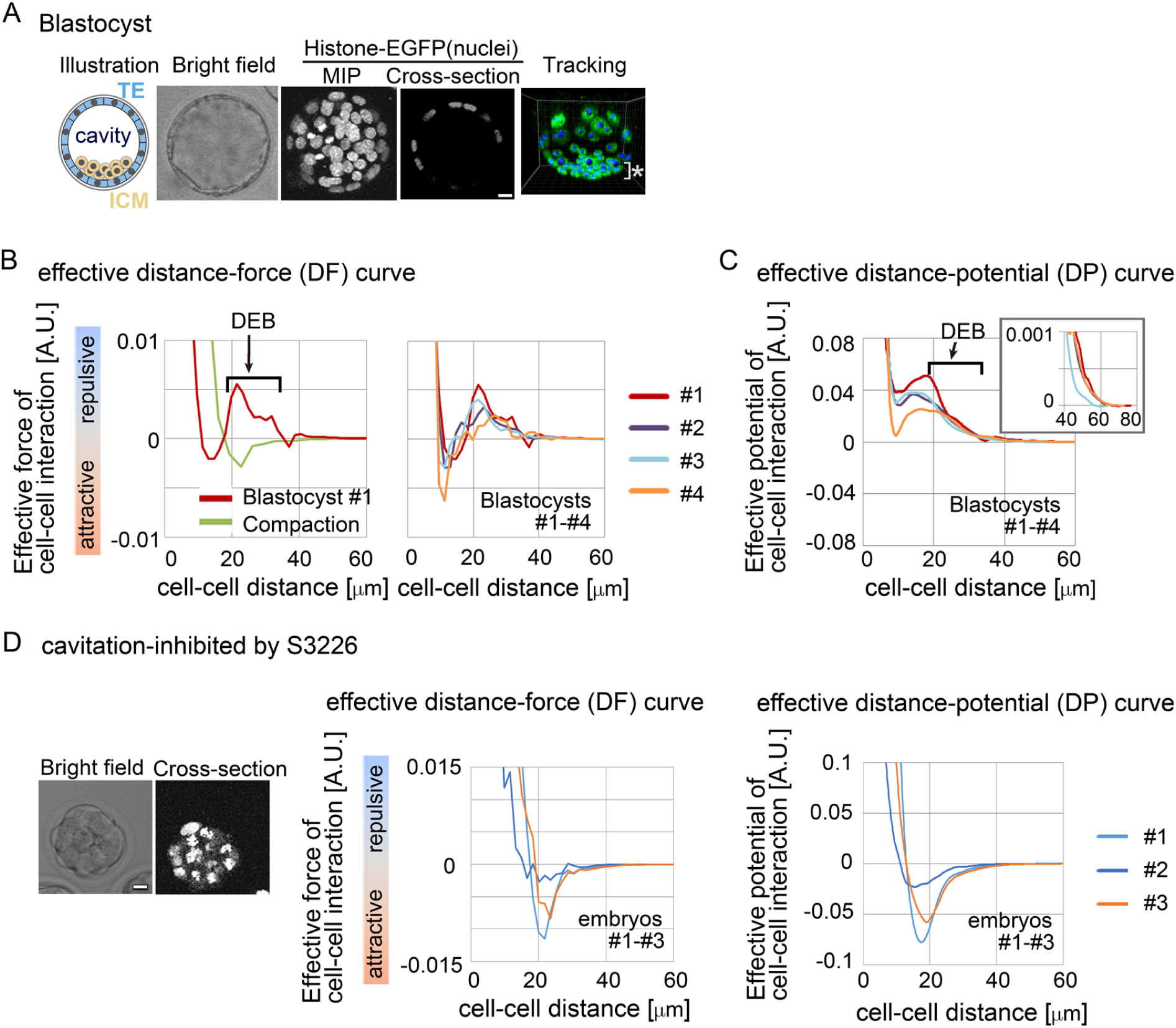
Inference of the effective force of cell–cell interaction in mouse blastocysts A. Blastocyst stages of mouse embryo are illustrated, and their confocal microscopic images are shown: bright field, maximum intensity projection (MIP) and cross-section (equatorial plane) of fluorescence of H2B-EGFP. Snapshots of nuclear tracking are also shown; blue spheres indicate the detected nuclei. ICM cells are located around the bottom region of the image, as indicated by *. Scale bars = 15μm. B. Inferred effective DF curve in the mouse blastocysts. In left panel, the blastocyst (#1) is shown by a red line. DEB is indicated. For comparison, an inferred effective DF curve from the compaction stage embryo before forming cavities is shown which we previously reported (green line) (Koyama et al., 2023). In the right panel, inferred effective DF curves from 4 blastocysts (#1-#4) are shown, whose bright field (BF) images are shown in Fig. S2A. C. Inferred effective DP curves in the mouse blastocysts. Four embryos were examined. An enlarged view around longer distance is also shown in the inset. The cell numbers in the 4 embryos and their changes during imaging were as follows; 56->58, 68->76, 56->65, and 58->59. D. Inferred effective DP curves in the mouse embryos where the cavitation was inhibited. S3226 whose target is Na+/H+ exchanger inhibits the cavitation of the blastocyst (Kawagishi et al., 2004). In the left panel, the microscopic images are shown. In the middle panel, the DF curves from three embryos are shown; the light blue line is from the embryo in the left panel. In the right panel, the DP curves are shown. The cell numbers in the 3 embryos were as follows; 32, 37, and 32 (the numbers were not changed during imaging). Scale bars = 15μm. Related figure: Figure S2 (inferred DF and DP curves based on Model-B) and S3 (experimental procedures of S3226 treatment)

Fig. 3B shows the inferred effective DF curve from the blastocyst. The DF curve contained the repulsive component around distant regions (Fig. 3B, DEB). For comparison, an inferred DF curve from an embryo before forming a cavity is shown, which did not contain DEB (Fig. 3B, “compaction” meaning the compacted embryo at the morula stage). In addition, Fig. 3C shows the distance–potential energy (DP) curves (*U*_C–C_(*D*)) which can be obtained by mathematically calculating the integral of the DF curves (*F*_C–C_(*D*)) along the distance (*D*) (Koyama et al., 2023); DP and DF curves are mutually convertible. In the DP curves, the DEB was expressed as a long slope with a negative gradient.

To further examine whether the DEB was derived from the cavity, we inhibited the cavitation by using an inhibitor for Na^+^/H^+^ exchanger (S3226) (Kawagishi et al., 2004). Fig. 3D shows the inhibitor-treated embryos; the experimental procedures are shown in Fig. S3. Fig. 3D shows the inferred effective DP curves, where DEB was diminished, indicating that the cavity yielded the DEBs.

We also evaluated a model dependency. We inferred effective forces under the assumption of Model-B. As shown in Fig. S2B-C, the inferred DF and DP curves were similar to those in Fig. 3B-C. Therefore, model choice did not significantly affect the inferred forces.

### 3.3 Effective potentials reproduced cavity-harboring structures

Effective DF/DP curves are the summation of the multiple components including the excluded volume effect, adhesive forces, and external factors, and are useful to study behaviors of systems in molecular and colloidal sciences. We evaluated whether inferred effective DF/DP curves from the cavity-harboring systems can reproduce their morphologies. The effective DF curves inferred from the mouse blastocyst, and the artificial tissue harboring a cavity (Fig. 2-3) were used for simulations, and the systems were relaxed to find stable states. When initial configurations of particles were set to be cavity-harboring structures (Fig. 4A), the structures were maintained (Fig. 4B-C, left panels). To examine whether the DEBs contribute to the formation of the cavities, we artificially eliminated the DEBs from the DF curves (i.e., the DF curves were truncated so as not to include the DEBs), and performed simulations. We found that cavities were not maintained but aggregates — filled with the particles — were formed (Fig. 4B-C, “DF curve w/o DEB”). For comparison, we used the Lennard-Jones potential, and showed that the cavity-harboring structure was not maintained but became an aggregate (Fig. 4D). In addition, when we used a DF curve inferred in cysts of Madin-Darby canine kidney cells, a cavity was also maintained (Fig. S4). In the case of the blastocysts, the particles were sparsely distributed (Fig. 4B). This was caused by the very long range of DEB in the blastocysts. In the blastocysts, the mean diameters (defined in Fig. 1C) were ∼10μm (Fig. 3B-C), and the ranges of DEBs were from ∼15μm to ∼60μm (Fig. 3C and its inset), meaning 1.5∼6.0-fold of the mean diameters. On the other hand, in the Lennard-Jones simulation data, the mean diameters were ∼5μm (Fig. 2C), and the ranges of DEBs were from ∼7μm to 15μm (Fig. 2A), meaning 1.4∼3.0-fold of the mean diameter. Note that these simulations explained the maintenance of cavities but not the generation of cavities from filled aggregates.

**Figure 4:**
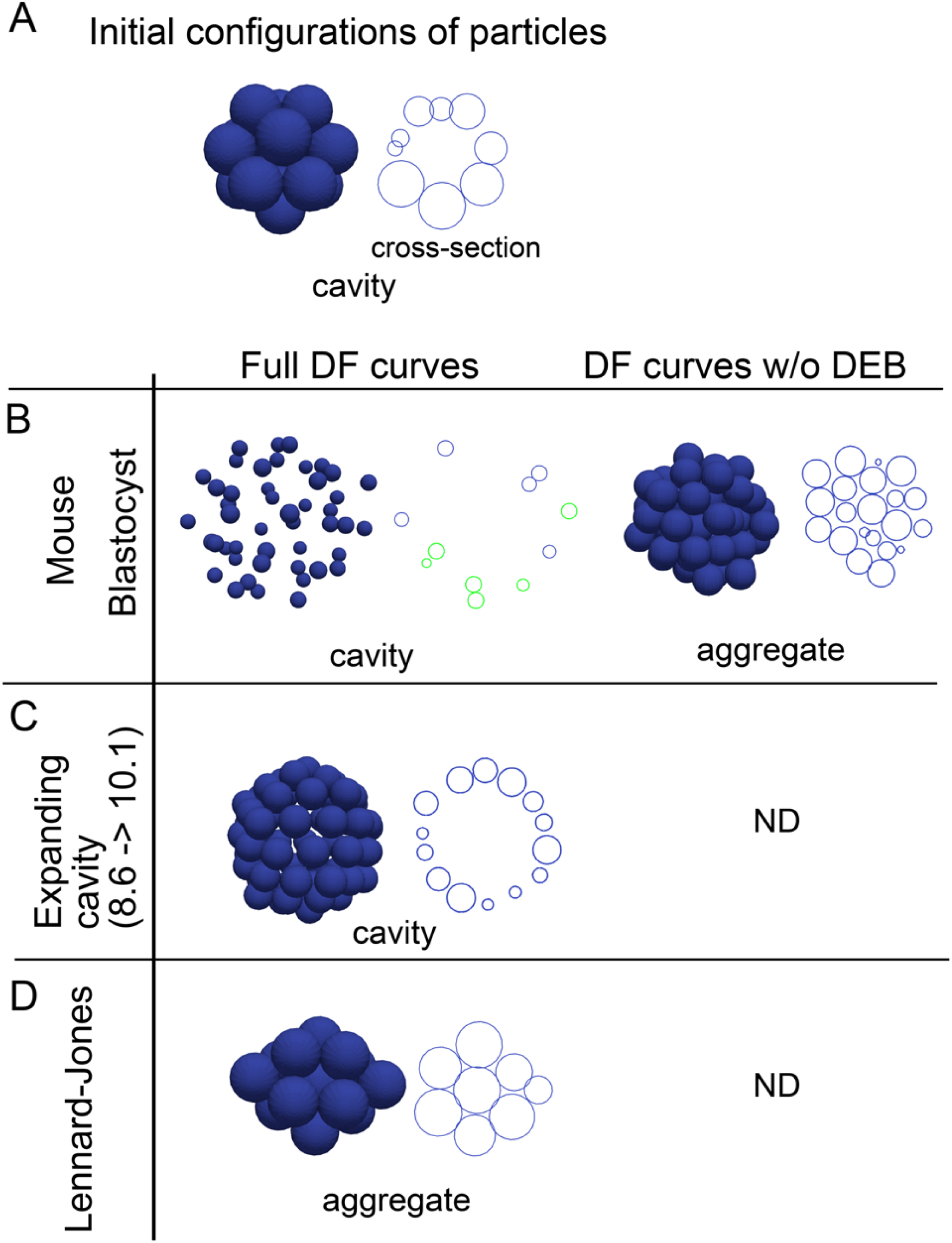
Outcomes of simulations based on inferred distance–force curves with DEBs A. Initial configurations of particles are exemplified, that harbor a cavity. A cross-section is also shown in the right panel. B-D. Simulations results under the effective DF curves derived from the mouse blastocyst (B), and the synthetic tissue harboring an expanding cavity (C) in Fig. 2 are shown. The DF curves used are from #1 in Fig. 3B for the blastocyst, and from Fig. 2A (8.6->10.1). E is the simulation result under the LJ potential for comparison. In addition to the left panels (“Full DF curves”), the DF curves in which the DEBs were artificially eliminated were used in the right panel (“DF curves w/o DEB); in C and D, analyzes for the right panel were not performed (“ND”). For each simulation result, cross-sections are also shown, and in the case of the mouse blastocyst, two parallel cross-sections (shown in blue and green) are merged. The diameters of the blue spheres are equivalent to the mean diameter.

### 3.4 Modeling of distance–force curves with two parameters

To comprehensively understand the effect of the differences of the profiles in the DF curves on morphogenesis, we modeled the DF curves as a simple mathematical equation. To analyze the contribution of DEBs, the frameworks based on the LJ and Morse potentials were not suitable, because they do not contain repulsive components around longer distances. Hence, we selected the following equation (Fig. 5A, left panel, blue line):

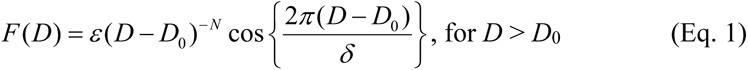

**Figure 5:**
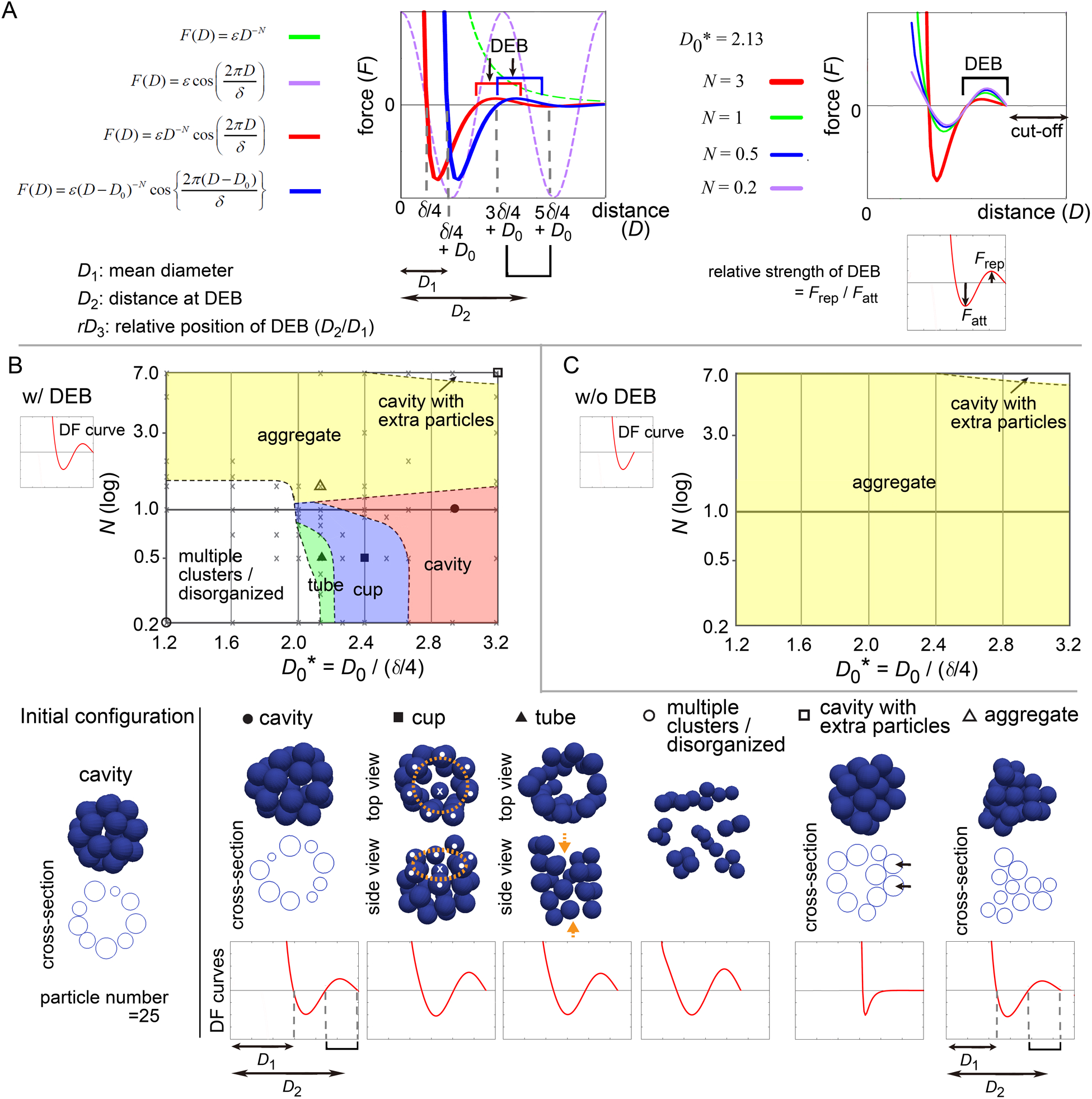
Modeling of distance–force curves and diagrams of simulation outcomes; cavity-harboring structures A. Modeling of DF curves using four parameters: *N*, *D*_0_, *δ*, and *ε*. In the left panel, a DF curve without the cosine term is always >0 (repulsive) (*F* = *εD ^−N^*, green broken line; *N* = 3, *ε* = 1.0). Upon introduction of the cosine (purple broken line), the curves acquire regions with <0 (attractive) and a DEB (red line; *N* = 3, *δ* = 1.5, *ε* = 0.2). Upon introduction of *D*_0_, the curves are translated along the distance (blue line; *D*_0_ = 0.3). As shown in the blue line, the mean diameter of particles corresponds to *δ*/4+*D*_0_ (*D*_1_), the DEB ranges from 3*δ*/4+*D*_0_ to 5*δ*/4+*D*_0_ where the center of the DEB is at *δ* + *D*_0_ (*D*_2_). In the right panel, *N* affects the gradient of the curves and the height of DEBs, and thus, the relative strength of DEB is modified. A cut-off distance was set: *F* (*D*) = 0, if *D* >5*δ*/4 + *D*_0_. B. Outcomes of simulations under various values of *N* and *D*_0_*\** in the presence of DEB. From the equation defined in A, DF curves from *D* = 0 to *D* at the end of DEBs (w/ DEB) were used as exemplified in the graph of DF curve (“w/ DEB”). The initial configuration is shown in the left bottom. On the diagram, some conditions were designated by symbols (circle, square, and triangle), and their simulation results are visualized on the bottom row (blue spheres) with the profiles of the DF curves. Cross-sections are visualized for some simulation results. The diameters of the blue spheres are equivalent to the mean diameter. For “cup” [orange rings, mouth of the cup; white dots, particles located on the ring; crosses, the particles located neither on the ring nor at the center of the ring]. For “tube” [orange arrows, open ends of the tube]. For “cavity with extra particles” [arrows, extra particles where multiple- but not single-layered]. The definition of each phase is explained in Supplementary information. Gray crosses, data points of all simulations for depicting the lines separating each phase (see Supplementary information). C. Simulation outcomes in the absence of DEBs. DF curves from *D* = 0 to *D* before starting DEB (w/o DEB) were used as exemplified in the graph of DF curve (“w/o DEB”). Related figures: Figure S5 (phase diagrams under a different number of particles), S6 (maintenance of tubes was examined under DF curves with DEBs), S7 (smoothness of the surface of the cavities), S8 (phase diagram for conditions of *D*_0_* < 1.2). Related movie: Movie S2 (the cavity-harboring structure was stably maintained under fluctuations)

Here, *F* is the force, *D* is the distance. *N* is the exponent of *D*, affecting the decay of the DF curves along *D* (X-axis) (Fig. 5A, left panel, green broken line). To model DEB, we introduced a cosine function whose wavelength is affected by *δ*; the combination of the above exponential function (Fig. 5A, left panel, green broken line) with the cosine function (Fig. 5A, left panel, purple broken line) resulted in a function bearing DEB (Fig. 5A, left panel, red line). We set a cut-off distance at the end of DEBs (i.e., *F* (*D*) = 0, if *D* >5*δ*/4 + *D*_0_ in Fig. 5A, right panel). *D*_0_ can translate the DF curves along *D*, (Fig. 5A, left panel, red vs. blue lines). By this translation, the mean diameter (*D*_1_) of particles is changed from *δ*/4 to *δ*/4 + *D*_0_, and the location of DEB becomes [3*δ*/4 + *D*_0_ ∼ 5*δ*/4 + *D*_0_] where the center of DEB is at the distance of *D*_2_ = *δ* + *D*_0_ (Fig. 5A, left panel). Therefore, this translation simultaneously modifies the relative positions (*rD*_3_) of DEBs to the mean diameters (*D*_1_) (Fig. 5A, left panel, *rD*_3_ = *D*_2_/ *D*_1_). *N* also affects the strengths of DEBs (Fig. 5A, right panel, *F*_rep_/*F*_att_ is the relative strength of DEB). *ε* determines the scale of the forces, that does not contribute to morphologies under stable states.

All parameters and variables related to lengths/distances were normalized by (*δ*/4), leading to generation of dimensionless parameters and variables such as *D*_0_*\** = *D*_0_/(*δ*/4). Therefore, we can present simulation outcomes in two-dimensional space (*D*_0_*\** and *N*) in a manner similar to a phase diagram (Fig. 5B) as shown in the next section. Although there can be other equations modeling DF curves, the advantage of this equation is that it is composed of just two parameters (*D*_0_*\** and *N*). All simulations were performed by using Model-A.

### 3.5 Appropriate position and strength of DEB were essential for cavity- harboring structures

First, we examined what kind of DF curves can maintain a cavity-harboring structures. In simulations of multicellular systems, the morphologies are determined by both DF curves and initial configurations, because energetic local minimum states (i.e. metastable state) as well as global minimum states are meaningful. A cavity-harboring structure was set as the initial configuration (Fig. 5B, “Initial configuration”), and simulations were performed until the systems reached steady states including metastable states; i.e., the velocities of the particles became almost undetectable.

Fig. 5B shows a phase diagram with representative images of the simulations, on which two regions of cavity-harboring structures were identified (“cavity” and “cavity with extra particles”). The “cavity” class absolutely depended on the DEBs, because DF curves whose DEBs were eliminated did not form the “cavity” as shown in Fig. 5C (“w/o DEB”; truncated version of DF curves exemplified at the upper left). We confirmed that the cavity was not broken by slight positional fluctuations (Movie S2 with the legend in Supplementary information), indicating that the structures were stable. In Fig. 5B, the “cavity” class was located around smaller *N* values (less than ∼2.0), otherwise the “aggregate” class was formed, suggesting that the strengths of the DEBs, defined in Fig. 5A (right panel), may be effective. Furthermore, larger *D*_0_ values were required for the “cavity” class, otherwise the “multiple clusters/disorganized” class was formed, suggesting that the relative positions of DEBs (*rD*_3_ = *D*_2_/ *D*_1_ in Fig. 5A, left panel) should be close to the mean diameters. Under a different number of particles, a similar trend was observed (Fig. S5; particle number = 64 and 150). The parameter values leading to the “cavity” class were almost equivalent between Fig. 5B and S5, and thus there was no clear relationship between the features of DEBs and the sizes of the cavities.

In addition, the “cups” and “tubes” classes were found between the “cavity” and the “multiple clusters/disorganized” classes in Fig. 5B but not in Fig. 5C, indicating that the DEBs also contribute to the formation of these structures. Because particle models are discretized systems, symmetry breaking for generating the cups and tubes can occur even under the directionally uniform DF curves. The maintenance of tubes was confirmed to be dependent on DEBs (Fig. S6). Detailed discussion about the “cup”, “tube”, “multiple clusters/disorganized”, and “cavity with extra particles” classes is described in Supplementary information.

The simulation outcomes of the “cavity” class showed slightly deformed spheres (Fig. 5B and S5A). We evaluated the smoothness of the surface of the cavity by calculating the distances of each particle from the center of the cavity (Fig. S7A). The values of the standard deviation (SD) of the distances were 3∼8% of the radii of the cavities (Fig. S7B-C). Therefore, absolutely smooth cavities were not generated by the DF curves with DEBs, which is a limitation of DEBs for describing spherical cavities. The SD values depended on both *D*_0_ and *N*; smaller values of *D*_0_ and *N* resulted in smooth cavities (Fig. S7B-C).

### 3.6 Profiles of DF curves for describing 2D sheet

To further investigate the capability of DF curves, we next applied a two-dimensional (2D) sheet as the initial configuration which corresponds to a monolayered cell sheet (Fig. 6A, “Initial configuration”). Fig. 6A shows a phase diagram with representative simulation images. The 2D sheets were maintained (the “sheet” class), which was largely dependent on the presence of DEBs (Fig. 6A, w/ DEBs vs. 6B, w/o DEBs; truncated version of DF curves exemplified at the upper left in Fig. 6B). We confirmed that the 2D sheets were not broken by slight positional fluctuations (Movie S2 with the legend in Supplementary information), indicating that the structures were stable. The region of the “sheet” class was located around larger values of *D*_0_, which was a similar trend to the case of the cavity in Fig. 5. The relationship between the sheet and cavity classes will be analyzed in detail later.

**Figure 6:**
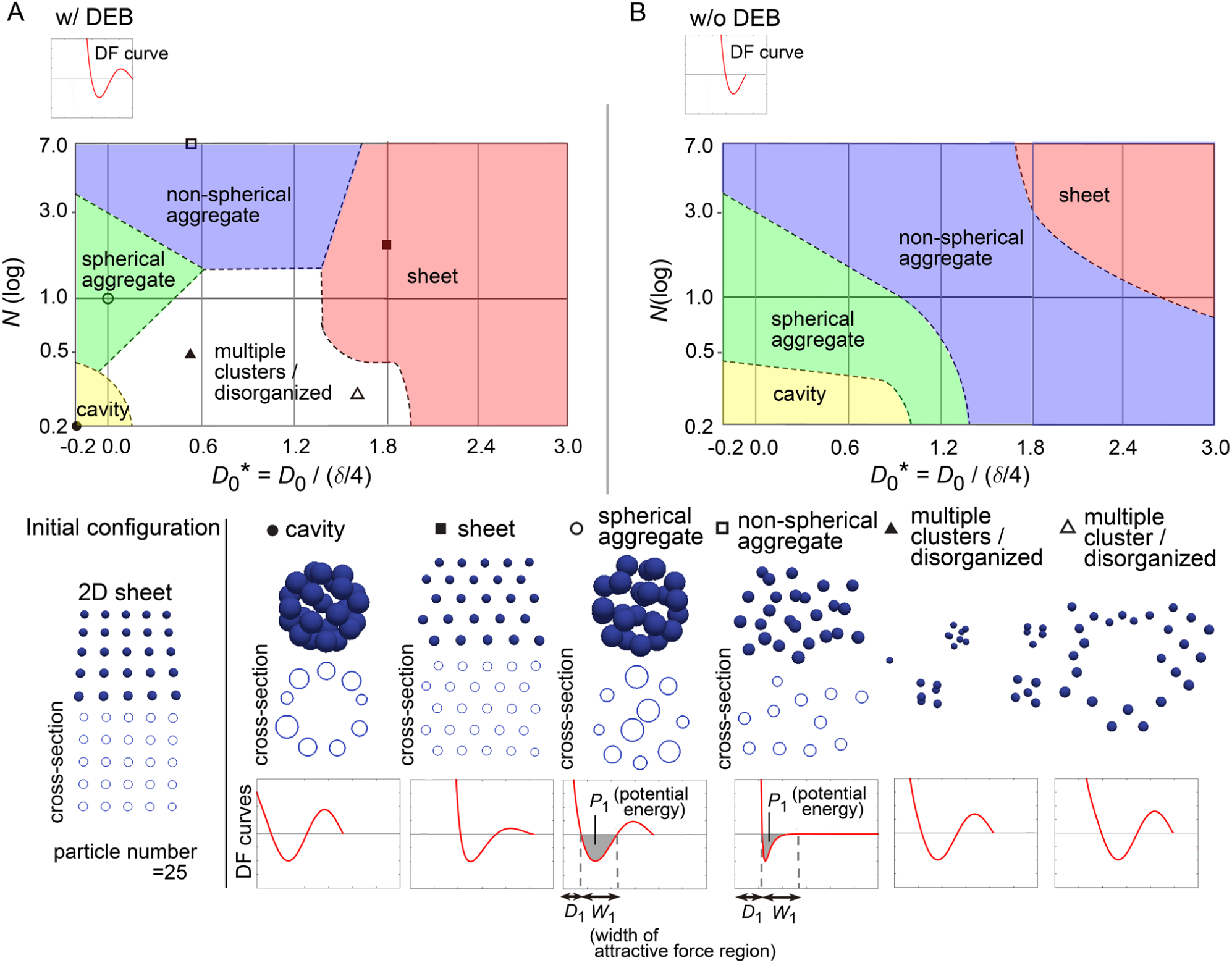
Diagrams of simulation outcomes; 2D sheets, spherical/non-spherical aggregates A. Outcomes of simulations starting from a two-dimensional (2D) sheet with slight positional fluctuations. The results are shown in a similar manner to Fig. 5B, except that the diameters of the blue spheres are equivalent to 0.4× the mean diameter. *D*_1_, mean diameter; *W*_1_, width of attractive force regions; *P*_1_, potential energy derived from attractive forces. Note that *D*_0_* can have a minus sign; because the mean diameter (*D*_1_) of a particle is *δ*/4 + *D*_0_, *D*_0_ should be > -*δ*/4 where *D*_0_* becomes > -1. B. Simulation outcomes without DEBs. Both Fig. 6B and 5C were generated under the conditions of w/o DEBs, but the ranges of *D*_0_* were different each other, resulting in that the “cavity” class was not appeared in Fig. 5C. We did not divide the “aggregate” class in Fig. 5C into the two “aggregate” classes as defined in Fig. 6B. Related figure: Figure S9 (possible morphological transitions) Related movie: Movie S2 (2D sheets were stably maintained under fluctuations), and Movie S3 (possible morphological transitions).

In addition, by expanding the parameter range in the phase diagram (i.e, *D*_0_* = - 0.2∼1.2), the cavity-harboring structures were newly generated even in the absence of the DEBs (Fig. 6B); because the parameter values were different from those for the cavity class in the previous figure (Fig. 5B), the cavity classes in Fig. 5B and 6B were different each other. Then, by applying the similar parameter values to the phase diagram in Fig. 5B, the cavity-harboring structures were also emerged (Fig. S8), indicating that the formation of this cavity class does not depend on initial configurations. Although we do not understand the precise mechanism of the de novo formation of the cavity, we guess that, if particles have a property to strongly assemble each other (e.g., due to strong cell– cell adhesion), the formation of a small cavity at the center can be energetically favorable; note that the sizes of the particles in Fig. 6B were set to be 0.4× mean diameter of the particles, and thus, the size of the cavity should be quite small. By temporally changing *N*, *D*_0_, and existence of DEB, we presented possible morphological transitions among the classes in Fig. 5 and 6 (Fig. S9 and movie S3).

### 3.7 Profiles of DF curves for describing spherical/non-spherical aggregates

In Fig. 6, we found that both the “spherical” and “non-spherical aggregate” classes were generated w/ and w/o DEBs. Smaller values of *N* and/or *D*_0_ led to the spherical ones (Fig. 6B). Under smaller values of *D*_0_, because the width (*W*_1_ ; see the bottom row in Fig. 6) of the attractive force regions in DF curves became relatively long compared to the mean diameter (*D*_1_, in Fig. 6), the resultant increased attractive forces may cause the spherical aggregates. Under smaller values of *N*, the potential energies (*P*_1_, in Fig. 6) derived from the attractive forces are increased, possibly leading to the spherical aggregates.

### 3.8 Effective forces in 2D sheets included DEB

As shown in Fig. 6, 2D sheets were maintained under the DF curves including DEB. Similarly, the cups and tubes in Fig. 5 were also maintained under the DF curves including DEBs; these structures can be interpreted as a curved sheet. As we previously showed, DEB was detected in tissues with pressurized cavities (Fig. 2-3). But it is unlikely that 2D sheets, cups and tubes have such pressures. Is the relationship between these structures and DEB an artifact or are there any reasonable explanations?

To investigate this issue, we used synthetic 2D sheets generated by simulations (Fig. 7A-i). The 2D simulations were performed under the Lennard-Jones (LJ) potential as the pregiven force, and the DF curves were inferred. As shown in Fig. 7A-i, the inferred DF curve was similar to the LJ potential, but the curve contained a clear DEB. This was contrast to the inferred DF curve from the simulation of a 3D aggregate, where DEB was rarely detected (Fig. 7A-ii).

**Figure 7:**
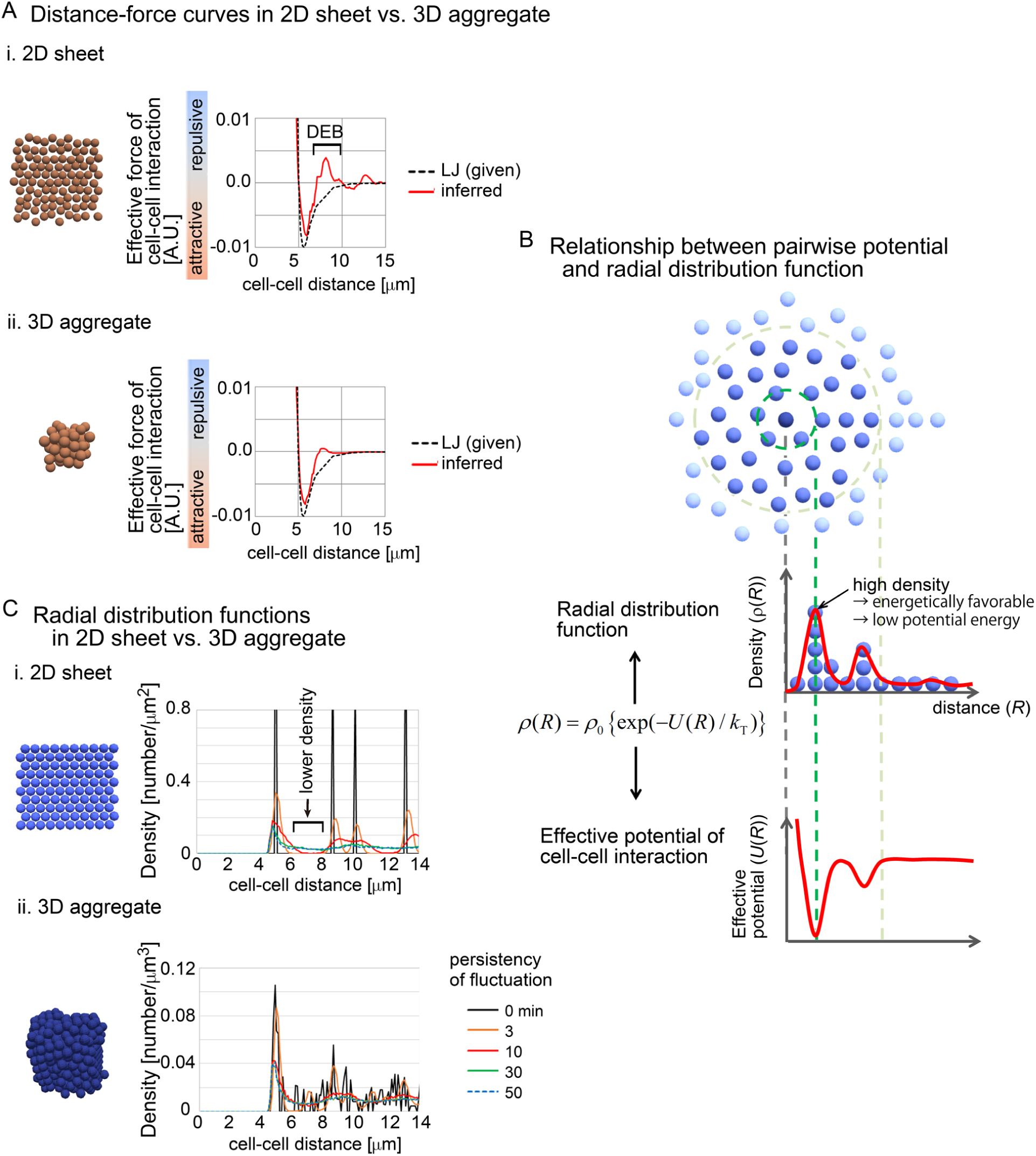
Radial distribution functions (RDF) in 2D and 3D situations A. Comparison of inferred DF curves in 2D (i) and 3D (ii) situations. The DF curves are shown with snapshots of simulations and with the pregiven potentials (LJ). DEB is indicated. The condition of the simulations is [sampling interval = 3 min, SD value of force fluctuation = 1,000%, and persistency of force fluctuation = 1 min]. B. The relationship between RDF and effective pairwise potentials is described. An RDF for the central particle (dark blue) is illustrated. Densities of particles are shown as the function of distances from the central particle. Higher density at a distance means an energetically favorable state, and therefore, the potential is expected to be lower at the distance. *k*_T_ is the effective temperature as an indicator of non-equilibrium fluctuations in multicellular systems, in analogy to *k*_B_*T* where *k*_B_ is the Boltzmann constant and *T* is the temperature. C. RDFs in 2D and 3D situations were calculated in simulations under the LJ potential. The snapshots of the simulations and the RDFs are shown. In these simulations, fluctuations of the forces given by the LJ potential were introduced with the persistency. Under longer persistency, the particles tend to easily change their positions. The condition is [SD value of force fluctuation = 300%, and persistency of force fluctuation = 1 min, sampling interval = 500min]. The numbers of particles are 100 (2D) and 500 (3D). Related figure: Figure S10 (distribution of particles under the closest packing situations under 2D and 3D conditions).

We hypothesized that the radial distribution functions (RDF) between 2D and 3D situations were different, because RDFs are related to the profile of distance–potential curves in atomic/molecular/colloidal sciences (Israelachvili, 2011). RDFs mean the density of particles at a certain distance from a particle of interest (Fig. 7B). The density may be reflected as the effective potential energy at the distance; high density corresponds to low potential energy (Fig. 7B). In other words, by using RDFs, effective pairwise potentials are calculated if the systems are under equilibrium conditions.

We calculated RDFs in 2D and 3D simulations performed under the LJ potential (Fig. 7C). In these simulations, fluctuations of the forces of cell–cell interactions were assumed with the persistency of the fluctuations (the details are described in Supplementary information). Under the 2D situations, clear peaks were found where the particle density was high. Importantly, between the peaks, the density was very low (Fig. 7C-i, black arrow). By contrast, under the 3D simulations, relatively unclear peaks were detected, and the regions with low particle densities almost disappeared (Fig. 7C-ii). These results may be reasonably interpreted by considering the closest packing situations under 2D and 3D conditions, where the peaks of high densities were frequently detected along the distances under the 3D condition compared with 2D (Fig. S10). Additionally, even under the 2D simulations, the longer persistency caused disappearance of the regions with low particle densities, because the long persistency promotes the movements of the particles. These results suggest that 2D sheets spontaneously yield regions with low particle density in RDF, probably due to the geometric reasons, and that this feature can be reflected to the effective pairwise potentials as DEB.

### 3.9 Expanding cavity was found before forming multiple clusters

We have been presented the two “cavity” classes in the phase diagram (Fig. 5B and 6). The formation of “cavity” class in Fig. 6 did not depend on DEB, and the “cavity” class in Fig. 5B may be derived from the geometric reason of cell sheets (i.e., curved sheets) because the location of the “cavity” class in Fig. 5B overlapped that of the “sheet” class in Fig. 6. In the case of the simulation of the blastocyst (Fig. 3B), the cavity was largely expanded, resulting in the sparse distribution of the particles. We failed to find such sparsely distributed pattern in the phase diagrams in Fig. 5, 6, and S5. Then, we carefully observe the dynamics of the simulations in Fig.5 and S5 whose initial configurations were “cavity”, and we found that, before forming the “multiple-clusters/disorganized” class, expanding cavities really existed. Fig. 8 shows the temporal dynamics of the simulations in Fig. S5 with the particle number = 64 (*D*_0_* = 1.33 and *N* = 0.2). The initial cavity was rapidly expanded, while maintaining its spherical shape. After it stopped expansion, the multiple clusters were formed. Therefore, the process of the expansion can be simulated by considering the DEB. Note that the boundary of the two origins of DEB may correspond to *D*_0_* ∼ 2.0, around which there is a risk to mix the two origins.

**Figure 8:**
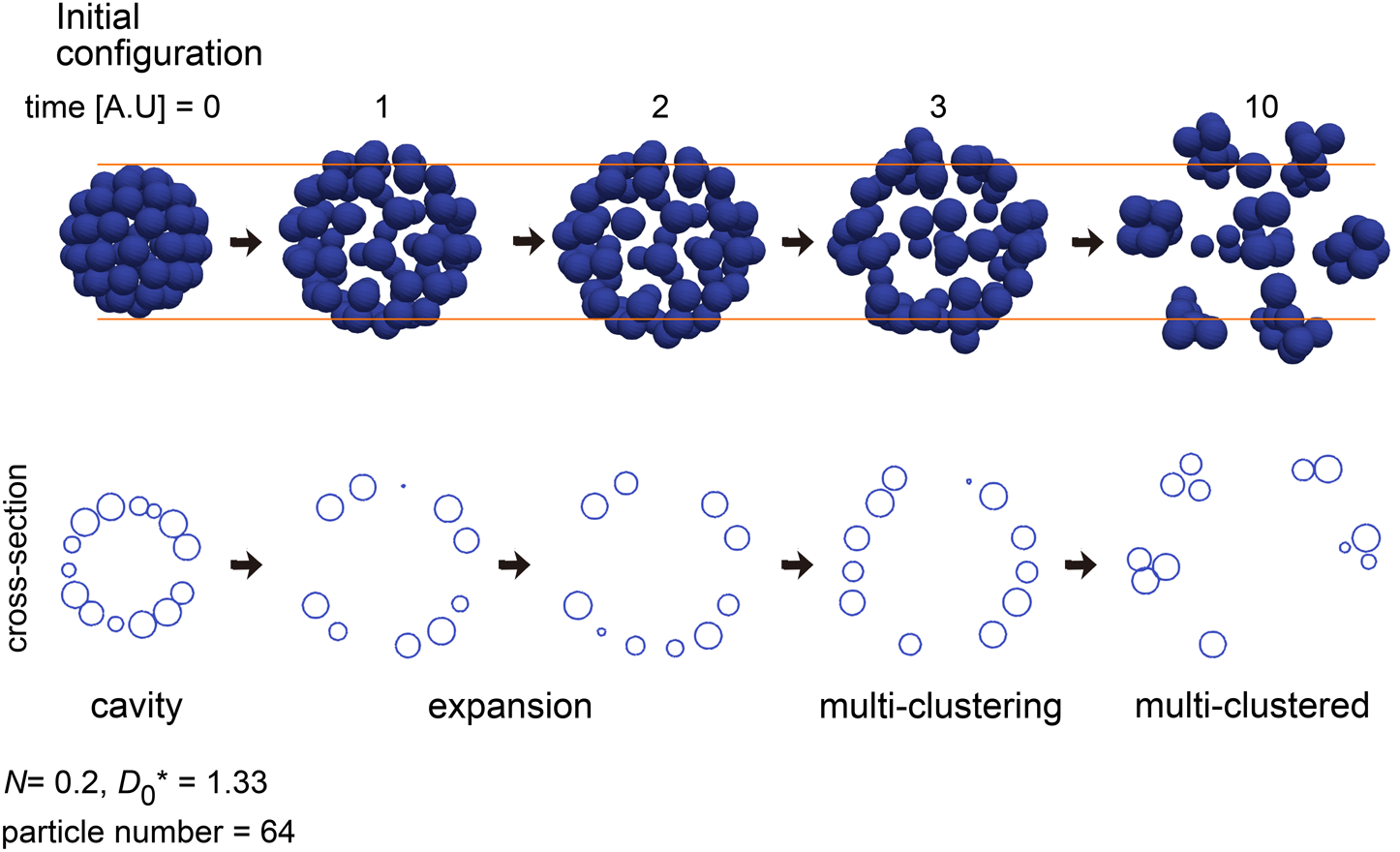
Simulation of expanding cavity An outcome of a simulation under a condition of *N* = 0.2 and *D*_0_*\** = 1.333 is shown with its temporal dynamics. Before forming multiple-clusters, the cavity was expanded. The particle number and simulation condition were the same as that in Fig. S5A.

## Discussion

In this study, by inferring effective forces of cell–cell interactions in tissues harboring expanding cavities, we showed that the effect of the cavities was addible to the effective forces of cell–cell interactions as a repulsive component (DEB). The resultant distance–force curves reproduced the tissue structures harboring cavities in simulations. A similar repulsive component was also detected in two-dimension (2D) cell sheets, which may be derived from geometric reasons of 2D systems. Finally, by assuming distance–force curves with the repulsive components, tissue structures of 2D sheets, cups, and tubes were reproduced in simulations; it is conceivable that, because 2D sheets, cups, and tubes may not harbor pressurized cavities, the origin of their DEBs is the geometric reasons of 2D or quasi-2D systems. These results suggest that external components and geometric constraints can be correctly added to the effective forces of cell–cell interactions. This framework considering the repulsive components may be expanded for simulating cell sheets with complicatedly curved surfaces such as epithelial folding as well as cavity- harboring structures, cups, and tubes (Wang et al., 1990a; Montesano et al., 1991; Honda et al., 2008b; Eiraku et al., 2011; Solnica-krezel and Sepich, 2012; Belmonte et al., 2016; Koyama et al., 2016; Nissen et al., 2018; Rivron et al., 2018; Serra et al., 2019).

### 4.1 Influence of external factors on effective potentials

Our results suggest that expanding cavities and geometric features of 2D sheet can be incorporated into the effective pairwise potentials. Another external factor in the mouse blastocyst is the Zona Pellucida (ZP) surrounding the embryo. In the late blastocyst, the ZP is outwardly pushed by the pressure of the cavity through the TE cells when the blastocysts are inflating, resulting in that the ZP is stretched. Conversely, the stretched ZP inwardly pushes the TE cells. Therefore, we think that the ZP should effectively reduce the magnitude of the vector from the pressure in Fig. 2D, “Pressure from cavity (red vector)”, meaning that the effect of the ZP is also incorporated into the inferred effective potentials. In atomic/molecular/colloidal sciences, it is known that some external factors are additive to effective pairwise potentials as mentioned in the Result section; e.g., hydration of ions, external pressures which can modify RDF (Israelachvili, 2011).In multicellular systems, it is unknown what kind of external factors are additive to the effective pairwise potentials. We speculate that, if external factors uniformly exert forces on cells, the effect may be correctly incorporated into the effective potentials.

In our present study, we focused on tissues with a single cavity of relatively larger sizes. On the other hand, at the beginning of the cavitation in the mouse embryos, multiple cavities with sizes much smaller than the cell size are generated (Dumortier et al., 2019). Smaller cysts are also known in the MDCK cells and other tissues (Wang et al., 1990b; Jewett and Prekeris, 2018). In such cases, particle model-based descriptions do not have sufficient spatial resolutions to simulate their morphologies, whereas other high-resolution models including the phase field models are suitable (Nonomura, 2012). On the other hand, because such small cavities are generated on cell–cell junctions by partially breaking cell–cell adhesion, it might be possible that, in our inference method, the generation of small cavities could be reflected as repulsive components of cell–cell interactions.

In the blastocysts and MDCK cysts, hydrostatic pressure of the cavities largely contributes to the cavitations. On the other hand, there can be other type of cavitation/lumenogenesis where hydrostatic pressure may be lesser (Jewett and Prekeris, 2018; Asleh et al., 2023). We do not know whether DF curves in such tissues become similar to those in the blastocysts, the MDCK cysts, or “cavity class” in Fig. 6.

According to our result that the effect of external factors is additive to the effective potentials, it is speculated that, if one can assume that external factors are equivalent between two tissues (e.g., wild-type and mutant tissues), differentials of the inferred effective DF curves between the two tissues correspond to the differences of cell– cell interaction properties. Therefore, even in the case that the precise effect of the external factors is not experimentally measurable, one can evaluate the differences of cell–cell interaction properties which may involve cell–cell adhesion. Similarly, we can also predict what will happen if cell-cell interaction forces are modified under the condition of non-cellular components. For instance, under the assumption of the fixed condition of non-cellular components (e.g., fixed expansion rate of cavities), we can simulate and predict cell behaviors in the case that cell-cell interaction forces such as adhesive forces are modified.

### 4.2 Relationship of the effective potential to cell shape, polarities, temporal changes, and cell–cell junction tension

In our simulations (e.g., Fig. 5-6), neither cell shape nor polarities were explicitly considered. On the other hand, the TE cells in the blastocysts exhibit a highly stretched squamous epithelium and an apico-basal polarity (Latorre et al., 2018). In addition, polarized tissues can have the planar cell polarity (Koyama et al., 2019). Although a cell in particle models look like a spherical cell shape, cell shapes and polarities can be considered to some extent by assuming anisotropy of the pairwise potentials; i.e., direction-dependent profiles of the potentials (Nissen et al., 2018). Such anisotropic potentials can elevate the capability of describing complicated tissue morphologies (Nissen et al., 2018). From our inferred forces, we may extract anisotropic potentials in principle if the anisotropy exists; a potential perpendicular to apico-basal polarity and a potential along apico-basal polarity. In addition, the TE cells are gradually stretched during the expansion of the blastocysts, and thus, the distance–force curves may be temporally changed. In the case of epithelial cells, cell–cell junction tensions are often considered and measured (Ishihara and Sugimura, 2012). The relationship between the profiles of the effective potentials and the cell–cell junction tensions is not understood, and we can theoretically analyze it by applying our inference method to simulation outcomes generated by the epithelial cell models such as the vertex model where the cell– cell junction tensions are assumed.

### 4.3 Comparison of other inference methods with other models

To understand morphogenetic events, simulations based on mechanical parameters are essential. Models such as vertex, Cellular Potts, and phase field models are often utilized, especially for epithelial cells (Honda et al., 2008a; Nonomura, 2012; Camley et al., 2014; Fletcher et al., 2014; Belmonte et al., 2016). These models successfully recapitulated various morphogenetic events, including the formation of the mouse blastocyst (Honda et al., 2008a). In these models, sub-cellular structures such as cell–cell junctions and their line tensions containing adhesive forces are explicitly modeled. In particular, the vertex models recapitulated well the behaviors of epithelial cells exhibiting polygonal cell shapes and three-dimensional cells exhibiting polyhedrons (Okuda et al., 2015a). On the other hand, non-polygonal epithelial cells are also found where cell–cell junctions are curved or interdigitated (Ishimoto and Morishita, 2014; Miyazaki et al., 2023), and thus, corresponding simulation frameworks become required. Because particle-based models are coarse-grained models, they cannot explicitly express cell–cell junctions. Due to their lower costs for programing, particle models were applied to three-dimensional simulations including intricately curves cell sheets (Nissen et al., 2018). Moreover, particle models were also applicable to mesenchymal cells that should exhibit intricated shapes (Paramore et al., 2024).

For the simulation of multi-cell behavior, the acquisition of parameter values is important. However, it is still challenging to measure cellular and non-cellular parameters in three-dimensional systems with high spatiotemporal resolution. Image-based force inference methods are important techniques (Tambe et al., 2011; Ishihara and Sugimura, 2012). Our idea can reduce the degrees of freedom derived from non-cellular factors, and may be adapted to other image-based inference methods when non-cellular factors exist. In analogy to the discussion in 4.1, differentials of forces between two tissues may be evaluated in the above image-based force inference methods. Our goal is to develop a framework for explaining diverse morphogenesis in a minimal model. Approximations of some external factors as effective potentials of cell–cell interaction serve to reduce degrees of freedom.

## Materials and Methods

### 5.1 Mouse embryos

Embryos at the embryonic days 2.5 (E2.5) were obtained after mating homozygous R26- H2B-EGFP knock-in male mice which constitutively express EGFP (enhanced green fluorescent protein)-fused H2B (histone2B proteins) (Kurotaki et al., 2007) and ICR female mice (Japan SLC). The embryos were cultured in EmbryoMax KSOM +AA with D-Glucose (Millipore, USA) covered with mineral oil on a glass bottom dish (35-mm; 27-mm ϕ, Matsunami, Japan) at 37°C. Then, the E4.0 embryos were subjected to microscopic imaging. Four distinct embryos were analyzed. Note that the mouse blastocysts sometimes lose the integrity of the sealing conferred by the tight junctions in TE cells, resulting in reduction of the pressure of the cavities with the subsequent volume decrease of the cavities (Niimura, 2003; Chan et al., 2019). This event is called “collapse”. Our four embryos, however, did not experience the collapse events during our microscopic imaging.

In the experiments using S3226 (Fig. S3), the E2.5 embryos were transferred to the medium with the drug. After 15 hours culture, some embryos failed to form cavities (Fig. S3, #1 and #2), while others formed smaller cavities (Fig. S3, #3) than that under no-drug conditions (Fig. S3, No drugs). Live imaging was performed, and then a rescue process was carried out, resulting in that the embryos formed, at least, smaller cavities compared with that under no-drug conditions. The concentration of S3226 was 50μM. We also tried Ouabain (Na^+^/K^+^ ATPase inhibitor)(Chan et al., 2019), but the cavitation was not drastically inhibited, and thus the embryos were not suitable for the cavity (-) control.

Animal care and experiments were conducted in accordance with National Institutes of Natural Sciences (NINS), the Guidelines of Animal Experimentation. The animal experiments were approved by The Institutional Animal Care and Use Committee of NINS (ref. #17A030, #18A026, #19A021, #20A014, and #21A039).

### 5.2 Microscopic live imaging

Fluorescence images were acquired on a Nikon A1 laser scanning confocal microscope (Nikon, Japan) equipped with a 20× objective (PlanApo; Dry; NA=0.75, Nikon, Japan). A stage-top CO_2_ incubation system was used (INUG2-TIZ, Tokai Hit, Japan) on the inverted microscope. H2B-EGFP was excited using a 488-nm laser.

For imaging of mouse embryos, 50-100 frames were acquired at 3-min intervals in 30–40 z-slices separated by 2.5 µm.

The confocal images obtained from the mouse embryos were subjected to the procedure for nuclear detection and tracking. Using the Imaris software (Oxford instruments/Bitplane, UK), the nuclei were automatically detected, followed by manual corrections. The nuclear tracking was also performed automatically followed by manual corrections. Through the manual corrections, the accuracies of the nuclear detection and tracking became essentially 100%, to the best of our judgment.

### 5.3 Inference method and simulation procedures

The inference method is as we previously reported (Koyama et al., 2023), where handling of cell division and cell death were also defined. Simulation procedures are also as previously reported (Koyama et al., 2023), except that cavities are considered. Details of the modeling of the cavities are described in Supplementary information.

## Acknowledgments

We thank Drs. Hitoshi Niwa and Yayoi Toyooka for providing the piggy-bac plasmids and technical suggestions.

