## Supplementary information for "Effective mechanical potential of cell–cell interaction in tissues harboring cavity and in cell sheet toward morphogenesis"

### *Supplementary Material*

#### Contents


1. **Supplementary text**
    - 1.1. **Particle-based cell model**
    - 1.2. **Generation of synthetic data by simulations**
        - **1.2.1.** Force fluctuation and its persistency
        - **1.2.2.** Simulation of cavity-harboring systems
        - **1.2.3.** Definition of heat map
    - 1.3. **Simulation based on inferred distance-force curves**
    - 1.4. **Modelling of distance-force curves and phase diagrams**
        - **1.4.1.** Modelling of distance–force curves
        - **1.4.2.** Phase diagrams
    - 1.5. **Possible transitions among various 2D and 3D structures achieved by modeled distance-force curves**
    - 1.6. **Experiments of MDCK cells**
2. **Supplementary Figures, Videos, and Sources**

### 1. Supplementary text

#### 1.1. Particle-based cell model

In our particle-based cell model, pairwise interactions of two particles are considered. The equation of particle motion is defined as follows: $V_{C|p} = F_{C|p} / \gamma$ (Eq. S1), where $V_{C|p}$ is the velocity of a particle, $p$ is an identifier for particles, $F_{C|p}$ is the summation of cell–cell interaction forces exerted on the $p$th particle, and $\gamma$ is the coefficient of viscous drag and frictional forces. We assumed the simplest situation, i.e., $\gamma$ is constant (=1.0) as described previously (Koyama et al., 2023). The simulations using distance–force curves were performed by the Euler method written in C.

In our inference method, a following cost function is minimized as described previously (Koyama et al., 2023); $G = G_{xyz} + G_{F_0}$. The first term ($G_{xyz}$) is the difference between $xyz$-coordinates of each particle of *in vivo* data and particle simulations as follows;

$$G_{xyz} = \sum_{t=1}^{T} \sum_{p=1}^{P(t)} \frac{\left\{x_p(t) - x_p^{\mathrm{ref}}(t)\right\}^2 + \left\{y_p(t) - y_p^{\mathrm{ref}}(t)\right\}^2 + \left\{z_p(t) - z_p^{\mathrm{ref}}(t)\right\}^2}{\Delta t},$$

where $p$ is the particle ID, $P(t)$ is the total number of particles at $t$, and $\Delta t$ is the time interval. $x_p$, $y_p$, and $z_p$ are the $xyz$-coordinates of $p$th particles in the particle simulations. $x_p^{\mathrm{ref}}$, $y_p^{\mathrm{ref}}$, and $z_p^{\mathrm{ref}}$ are the $xyz$-coordinates of $p$th particles in the *in vivo* cells. $G_{F_0}$ is the additional cost function for cell–cell distance-dependent weight as previously defined (Koyama et al., 2023); $G_{F_0} = \omega_{F_0} \sum_{t=1}^{T} \sum_{i=1}^{I(t)} \psi(D_i) F_i^2 \Delta t$, where $F_i$ is the force value of $i$th cell–cell interactions, $I(t)$ is the total number of the cell–cell interactions for each time frame, $D_i$ is the cell–cell distance of the $i$th interaction, and $\omega_{F0}$ is a coefficient for determining the relative contribution of the second term to $G$. $\psi(D_i)$ is a distance-dependent weight that exponentially increases and eventually becomes extremely large, leading to that the value of $F_i$ approaches zero especially around larger $D_i$.

#### 1.2. Generation of synthetic data by simulations

##### 1.2.1. Force fluctuation and its persistency

We generated simulation data which were used for validating our inference method as described previously (Koyama et al., 2023). Briefly, these simulations were performed under the distance–force (DF) curves derived from the Lenard-Jones potential, which can be written as follows;

$F(D)=\frac{\kappa}{\sigma}\left\{12\left(\frac{\sigma}{D}\right)^{13}-6\left(\frac{\sigma}{D}\right)^{7}\right\}$ as a DF curve. $\sigma$ corresponds to the distance where the potential becomes 0, and $\kappa$ determines the scale of the forces. In addition, we introduced fluctuations of the cell–cell interaction forces as relative values of the forces derived from the DF curve as follow: $F_{\mathrm{wfl}}(D) = F(D) + v\,(pc,\,\tau)\,|F(D)|$, where $F_{\mathrm{wfl}}$ is the force of cell–cell interactions, wfl means "with fluctuation", and $v$ is the magnitude of the fluctuations defined as a ratio to $|F(D)|$. $v$ was defined from minus to plus values. $v$ is given by Gaussian distribution, $N(0, (pc/100)^2)$, where the average and standard deviation (SD) of the Gaussian distribution is 0 and ($pc/100$), respectively. $pc$ means the percentage. $\tau$ is the persistence time period reflecting time scale of cellular processes. The value of $v(pc,\tau)$ is evolved for every time period $\tau$ and is unchanged during $\tau$. In the related figures, $pc$ and $\tau$ are shown as "SD value of force fluctuation (%)" and "Persistency of force fluctuation (min)", respectively. The precise definition was described in our previous work (Koyama et al., 2023).

##### 1.2.2. Simulation of cavity-harboring systems

For the simulations of cavity-harboring systems, in addition to cell–cell interaction forces defined by the LJ potential, a spatial constraint was implemented as a cavity. In Figures 2 and S1, when a particle moves to the inner region of the surface of the constraint, the position of the particle was modified outwardly from the center of the spherical constraint so that the position was just in contact with the surface (Fig. S1). On the other hand, particles can move outwardly from the surface of the constraint (i.e. there is no constraint for preventing the particles from dissociating from the surface of the constraint).

##### 1.2.3. Definition of heat map

The heat maps in Figure 2C were generated from the frequencies of the data points as defined in our previous report (Koyama et al., 2023); the graph space composed of the distance and the force was divided into 64×64 square regions, and then, the data points were allocated to one of the 64×64 square regions. We considered 64 columnar regions along the distance, and the mean values of the counts of the square regions were calculated for each columnar region. The mean count for each 64-column region was defined as the frequency index (FI) = 1. The lookup table of the colored map is shown in Figure 2C; many data points were plotted around the red regions, whereas few data points were plotted around the black regions. In the white regions, no data points were plotted.

#### 1.3. Simulation based on inferred distance–force curves

In Figure 4, cavity-harboring structures were provided as the initial configurations, and we tested whether the cavity-harboring structures were maintained after relaxation of the simulations. The numbers of particles were set to be different so that the numbers were nearly consistent with each *in vivo* situations (i.e. the blastocyst and MDCK cyst).

#### 1.4. Modelling of distance–force curves and phase diagrams

##### 1.4.1. Modelling of distance–force curves

We explored modeling of distance–force curves bearing distant energy barriers (DEBs). The LJ potential and its variants with different values of the multiplier factors cannot yield DEBs. To implement DEBs, we introduced the cosine function in an analogy to the RKKY (Ruderman–Kittel–Kasuya–Yosida) interaction as considered in metals. We considered as simple a function as possible, and defined Equation 1, $F(D) = \varepsilon(D - D_0)^{-N} \cos\left\{\frac{2\pi(D - D_0)}{\delta}\right\}$, in the main text, as one of the candidates. There are four parameters, $N$, $D_0$, $\varepsilon$, and $\delta$, which determine the profile of the distance–force curves. $\varepsilon$ determines the scale of the forces. Introduction of a dimensionless parameter $D_0^* = D_0/(\delta/4)$ enables us to produce a two-dimensional diagram of simulation outcomes (Fig. 5 and 6). Note that introduction of dimensionless parameter is a well-known technique in physics and theoretical biology, which does not alter simulation outcomes: under the same value of $D_0^*$, final morphologies of the simulations became the same even when different values of $D_0$ or $\delta$ were provided. $\delta/4$ refers to one-quarter of the length of a single wave of the cosine function. Therefore, when the distance is $\delta/4$, the force becomes 0 under the condition that $D_0 = 0$, meaning that the diameter of the core coincides with $\delta/4$. In the case of $D_0^* = 1$ ($D_0 = \delta/4$) as an example, the distance–force curve is translated along the distance by the length of the original core diameter; consequently, the core diameter is 2-fold greater than the original one.

##### 1.4.2. Phase diagrams

As in systems composed of larger colloidal particles which hardly reach a thermodynamically equilibrium state (Israelachvili, 2011), multicellular systems are usually in or near metastable states corresponding to local minima, or under non-equilibrium states; therefore, our focus was not limited to thermodynamically equilibrium states corresponding to global minima. To find stable states, including local and global minima, we started our simulations from various initial particle configurations. In other words, simulation outcomes depend on not only distance–force curves but also initial configurations.

In Figure 5, a cavity-bearing structure was used as the initial configuration of particles. We performed another analysis considering different particle numbers. In contrast to Figure 5, we found that tubes were not generated and cups were formed under limited conditions (Fig. S5). These results indicate that the probability of the emergence of tubes and cups was affected by the particle numbers. In Figure 6 where a two-dimensional sheet was applied as the initial configuration, slight positional noises were introduced for each particle in the initial configuration, because a mathematically absolute plane cannot deform along the direction vertical to the plane during simulations.

Definition of the phases, “cavity”, “cup”, and “tube”, in Figure 5, 6, and S5-7 is described as follows. The diameters of the particles in these simulations were defined to be the same as the distances at the potential minima of the DP curves (i.e. the mean diameter) used in the simulations (Fig. 5, S5-7) or 0.4× the mean diameter (Fig. 6). “Cavity”, “cup”, or “tube” has zero, one, or two holes on the surface of the simulation outcomes, respectively. Existence of holes was determined by calculating whether the space of interest on the surface is larger than a circle whose diameter is equivalent to the distance at the potential minimum in the DP curve.

The boundaries separating different phases on the phase diagrams were drawn according to the data points (Fig. 5B; data points were indicated by crosses). Basically, we manually drew the boundaries so that they passed between the crosses classified into different phases. Around intricate regions (e.g., Fig. 5B, around [$N$ = 1.0, $D_0$* = 2.0]), we increased data points. We also took a dependency of initial configurations of particles into consideration. Around some boundaries of different phases (e.g. the boundary between “cavity” and “cup” in Figure 5B), simulation outcomes were sensitive to the initial configurations of particles even under the same number of the particles with the same values of the parameters, $N$ and $D_0$. So, we performed three simulations with slightly different initial configurations that were generated by adding small positional noises for each particle; the noises were provided by uniform random numbers along x, y, and z axes with ± ~0.04% of the mean diameters of the particles. When the three simulation outcomes were different each other (e.g. one outcome is “cavity”, and other two outcomes are “cup” in Figure 5B), the boundary separating the two phases on the phase diagram was drawn just on the position of the parameter values used in the simulations.

The phases, “cavity” and “cavity with extra particles”, in Figure 5 were determined by evaluating whether the simulation outcome is solely composed of a single-layered cell sheet or not. The “cavity” is a single-layered structure, whereas the “cavity with extra particles” contains regions with multi-layered structures.

The “tube” in Figure S6C was determined in a manner independent to the sizes of the holes, but just by evaluating whether an empty space exist throughout the center line of the simulation outcome.

Among the “cavity”, “cup”, “tube”, and “multiple clusters/disorganized” classes, as $D_0$* is decreased, the relative positions ($rD_3$) of the DEBs become far from the mean diameters, implying that such far-repulsive forces may destabilize the structures, leading to the generation of multiple holes and eventually to multiple clusters. The “cavity with extra particles” class was found at larger $N$ (~7) where the repulsive forces of the DEBs are quite weaker, that is contrast to the “cavity” class. At the same time, the range of the attractive forces is narrow due to larger $D_0$*, meaning that the particles are not easy to be closely assembled to form the “aggregate” class.

#### 1.5. Possible transitions among various 2D and 3D structures achieved by modeled distance–force curves

By using the Equation 1, we investigated possible morphological transitions under the assumption of distance–force curves. We temporally changed the parameter values in the Equation 1 ($N$, $D_0$, and $\delta$) and with or without DEB. Figure S9 shows the morphological transitions which we were able to achieve. The transition contains; two-dimensional sheet → aggregate, two-dimensional sheet → cavity, aggregate → cavity, cavity → aggregate, cavity → cup, cup → aggregate, cup → tube, tube → aggregate, tube → cavity. Thus, DF curves are sufficient to describe transitions among 2D (2D sheet), quasi-3D (cavity, cup, and tube; i.e. curved sheets), and 3D (aggregate) structures. We can interpret that, if cells temporally experience such modulation of the profile, these modulations reflect temporal changes in properties of cell–cell interactions during morphogenesis.

#### 1.6. Experiment of MDCK cells

##### 1.6.1. Inference of effective forces

We applied our method to cysts of MDCK cells. MDCK cells can form cysts containing a cavity when cultured under suspension conditions (Fig. S4A) (Wang et al., 1990). Fig. S4B and S4C show the effective DF and DP curves. In the MDCK cysts, DEBs were detectable or hardly detectable (Fig. S4B-C). Under our culture condition, the cysts did not rapidly grow; it took 4-5 days to reach ~10 cell-cysts. Therefore, inflation of the cavities should be slower than that in the blastocysts. We think that, due to the slower inflation, DEBs were not so clearly detected. This is consistent with the theoretical prediction obtained from Fig. 2 and S1C.

##### 1.6.2. Construction of MDCK strain

MDCK cells were described previously (Otani et al., 2019), and were originally provided from Masayuki Murata (University of Tokyo). The identity of the cell line (derivation from canine) was confirmed by genomic sequencing of targeted loci. The mycoplasma was removed, and the removal was confirmed by Minerva biolabs Venor GeM OneStep Mycoplasma detection Kit for Conventional PCR.

PiggyBac transgenic MDCK cells which constitutively express H2B-EGFP were constructed as follows. To obtain a DNA fragment bearing H2B-EGFP, pcDNA3-H2B-EGFP (Abe et al., 2011) was digested with *HindIII*, blunted, and then digested with *NotI*. pEBMulti-Puro (Wako, Japan) was digested with *BamHI*, blunted, and then digested with *NotI*. Then, the DNA fragment bearing H2B-EGFP was inserted into the pEBMulti-Puro backbone. From this plasmid, a DNA fragment harboring H2B-EGFP was excised again by *XhoI* and *NotI* digestions, and was then inserted between the *XhoI* and *NotI* sites of pPB-CAG-BstXI-IP (a gift from Dr. Hitoshi Niwa) (Nakatake et al., 2013). The resultant plasmid, pPB-CAG-IP-H2B-EGFP, and the PBase-expressing vector pCAGPBase (Guo et al., 2009) were co-transfected into MDCK cells using Lipofectamine LTX (Invitrogen), and the transformants were selected in the presence of puromycin (1 μg/mL). The transformants were expected to express H2B-EGFP under control of the constitutive CAG promoter.

##### 1.6.3. Experimental procedure

To generate and culture cysts of MDCK cells harboring H2B-EGFP, MDCK-conditioned DMEM (Dulbecco's Modified Eagle's Medium containing 10% fetal bovine serum with NaHCO3, Sigma) was used. The conditioned medium was obtained as follows: MDCK cells were cultured in 10 mL DMEM on a 95-mm dish for 4 days at 37°C to reach ~90% confluence, 10 mL fresh DMEM was added, and the cells were cultured for an additional day. The supernatant was obtained, and filtered through a 0.22-μm Millipore filter (Millex-GV SLGV033RS, Millipore, USA), yielding conditioned medium.

The cysts were generated and cultured as follows: MDCK cell suspension in 80% MDCK-conditioned DMEM and 20% DMEM was placed on glass-bottom dishes coated with 0.4% agarose (SeaKem GTG agarose, BMA). The cells were cultured for 4–5 days at 37°C, and then subjected to microscopic imaging. Four distinct cysts were analyzed. For imaging of MDCK cysts, 50-100 frames were acquired at 10-min intervals in 20–30 z-slices separated by 2.0 μm.


#### 2. Supplementary Figures, Videos, and Sources

##### 2.1. Supplementary Figures

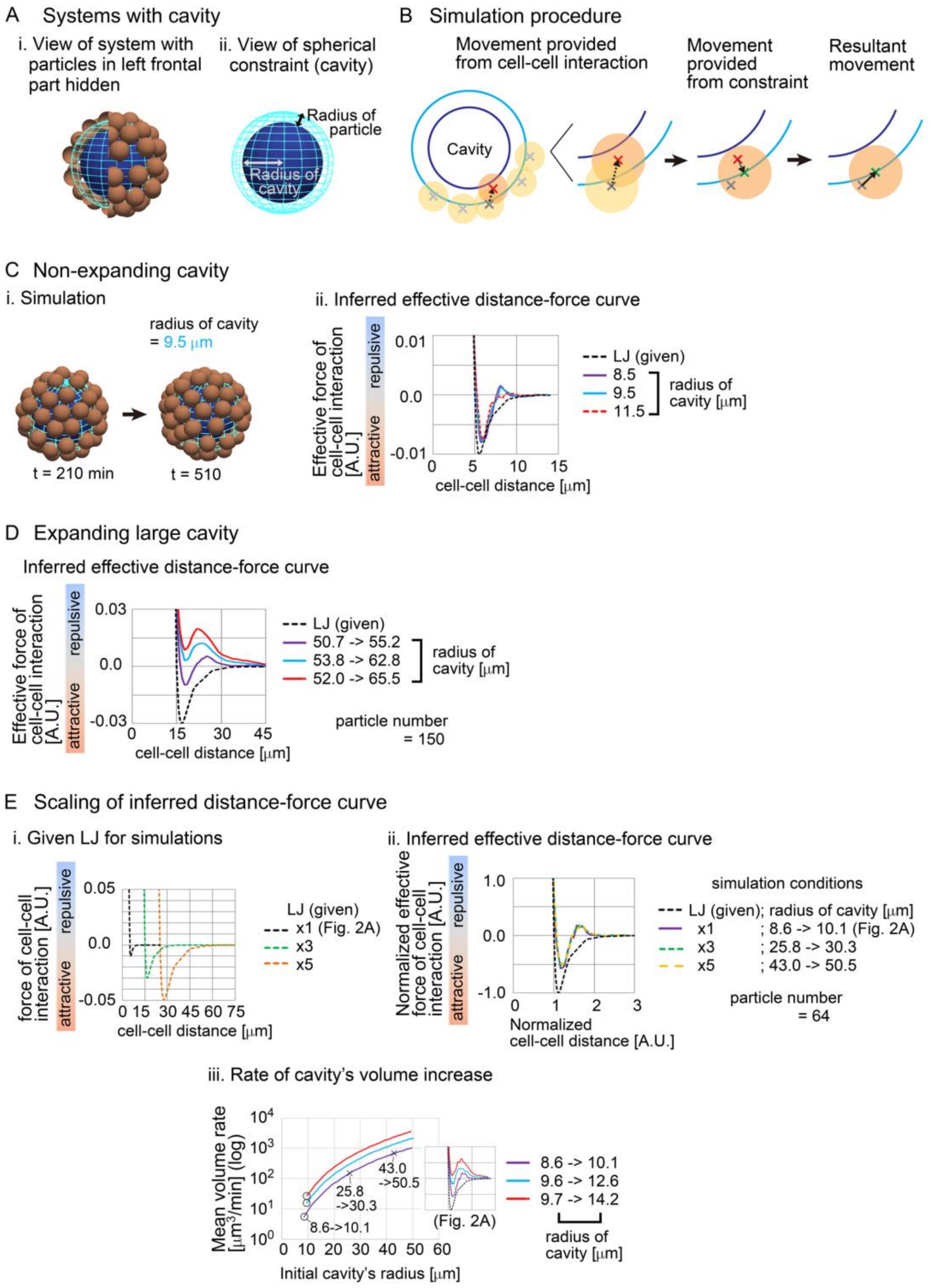


Figure S1 (related to Figure 2): Simulation procedures under cavities and inferred effective pairwise forces

A. Particles were put on the surface of spherical cavities (A-i). Definitions of the radius of the spherical cavity and the radius of particles are shown (A-ii).

B. The simulation procedures when a particle collides with the spherical cavity as a spatial constraint are described (dark orange circle). The detailed procedures are described in Supplementary information.

C. Inferred DF curves under different radius of cavities with snapshots of simulations. The cavities were not assumed to expand. The condition is [sampling interval = 3.0 min, SD value of force fluctuation = 1000%, and persistency of force fluctuation = 1 min]. The number of particles were set to be 64 in all conditions.

D. Inferred DF curves under larger radii of expanding cavities with a larger particle number (=150).

E. Scalability of inferred DF curves. i) Both the distances and forces from the LJ potential were magnified (×3 and ×5), which were used for generating simulation data. ii) Inferred DF curves in expanding cavities whose radii were set to be ×3 and ×5 compared to the "×1" condition which corresponds to Fig. 2A. In the case of the "×3" and "×5" conditions, the given DF curves were also set to be ×3 and ×5 as defined in (i). For comparison, the values of the inferred forces and distances were normalized so that the three LJ potentials became the same scale. Equivalent DF curves were obtained from the three conditions. iii) Rates of cavity's volume increase were calculated. In the case of the condition of radius = 8.6->10.1, the volume rate was from $10^0$ to $10^1$ marked by a black circle on the purple line. In the case that this condition was magnified as defined in (ii), the relationship between the initial radius and the volume rate is shown by the purple line; e.g., the conditions of "×3" and "×5" in (ii) are plotted by crosses. Similarly, other two conditions in Fig. 2A (radius = 9.6->12.6 and 9.7->14.2) correspond to the black circles on the light blue and red lines. The magnified results for the two conditions are shown by the light blue and red lines.

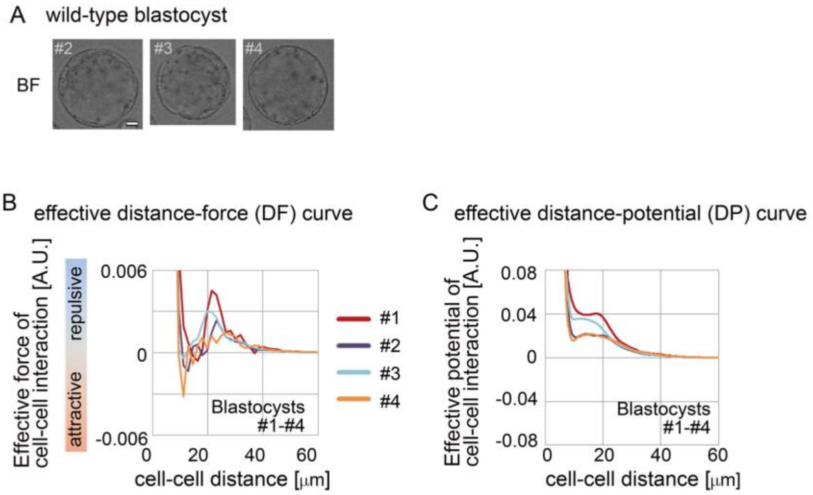


Figure S2 (related to Figure 3): Inferred effective pairwise force in blastocysts under Model-B
A. Bright field images of the blastocysts #2-#4 in Fig. 3. Scale bar, 15μm.
B. Inferred effective DF curve in the mouse blastocysts. The Model-B was used for the inference procedures as described previously (Koyama et al., 2023). $\gamma$ in Model-B was set 0.1, and $\omega_{\mathrm{g}}(D_{pm})$ was set as described previously. The blastocysts were the same as those in Fig. 3.
C. Inferred effective DP curves in the mouse blastocysts.

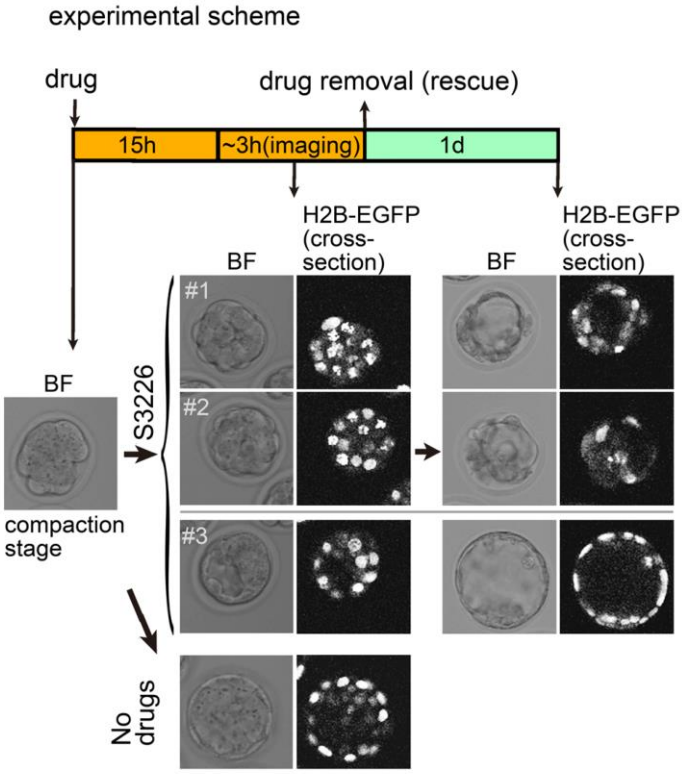


Figure S3 (related to Figure 3): Experimental procedure for cavitation-inhibited mouse embryos

The experimental design for inhibiting cavitation during the formation of the blastocysts are shown. The mouse embryos at the compaction stage were cultured in the presence of S3226, and the resultant embryos failed to form cavities (#1; the same image as that in Fig. 3D and #2) or formed a smaller cavity (#3) than the normal embryos (no drugs). After the removal of the drug (rescue), the drug-treated embryos formed cavities though the sizes of the cavities were often smaller (#1 and #2). Therefore, the drug-treatment was nonlethal for the cells. BF, bright field. Scale bar, 15μm.

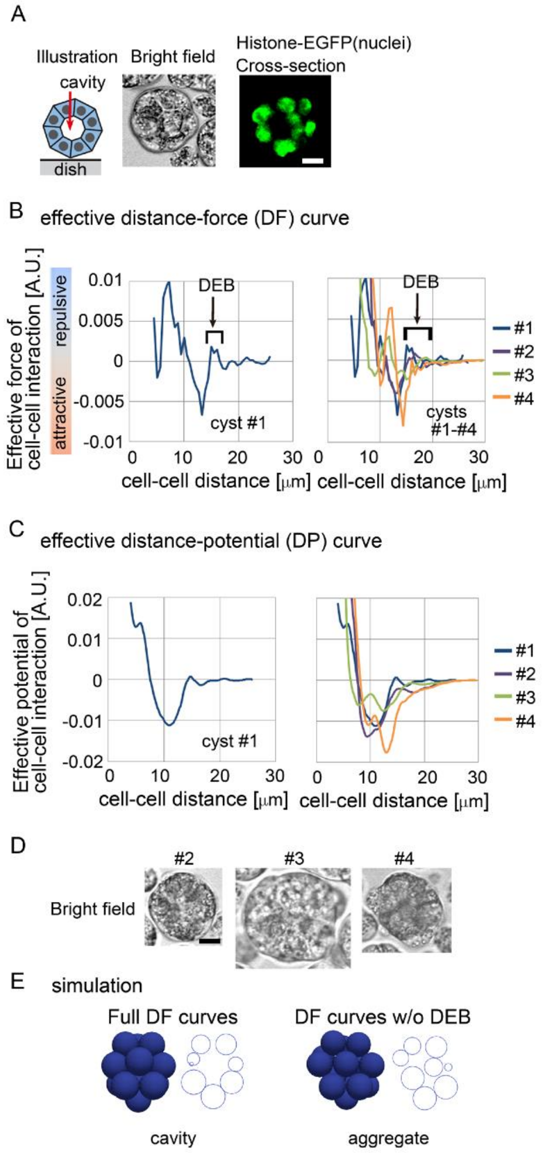


Figure S4 (related to Figure 4): Distance–force and distance–potential curves in MDCK cysts

A. MDCK cysts formed under suspension conditions are illustrated, and confocal microscopic images are shown: bright field, maximum intensity projection (MIP) and cross-section (equatorial plane) of fluorescence of H2B-EGFP. Scale bars = 10μm.

B and C. Inferred effective DF and DP curves. A DEB is indicated. Four independent cysts (#1-4) were analyzed. The cell numbers for the 4 cysts were as follows; 11, 12, 15 and 13 (the numbers were not changed during imaging).

D. The bright field (BF) images for the #2-#4 cysts are shown. Scale bars = 10μm.

E. Simulations results under the effective DF curves derived from the MDCK cyst #1 (Fig. S4B, left panel). Other simulation conditions and procedures are the same as those in Fig. 4.

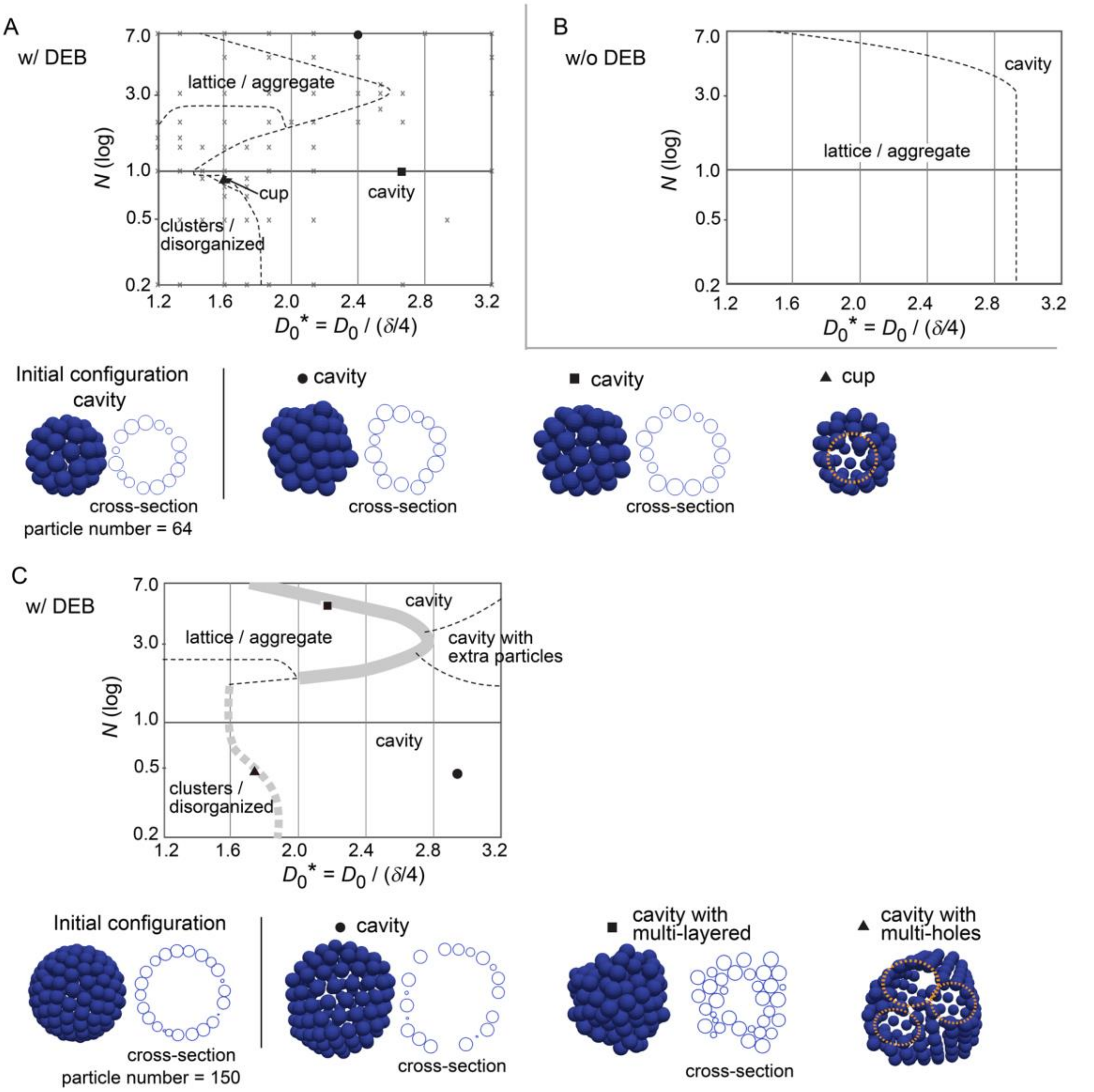


Figure S5 (related to Figure 5): Simulation outcomes and their diagrams, based on distance–force curves; starting from cavity-harboring structure

A-B. Phase diagrams are shown in a similar manner to those in Fig. 5, except that the numbers of the particles are different; particle number = 64. The relative location of the phases (e.g., "cavity") in the diagrams is almost similar to that in Fig. 5. The diameters of the blue spheres are equivalent to the mean diameter.

C. A phase diagram in the case of the particle numbers = 150 is shown in a similar manner to A. Around the boundary between the "cavity" and "lattice/aggregate" classes, multi-layered cavities were generated, resulting in that the boundary became obscure, which was shown by a bold gray line. Similarly, around the boundary between the "cavity" and "cluster/disorganized" classes, cavities with multiple holes including the "cup" class (single hole) were generated. This boundary became also obscure and was shown by a bold broken gray line.

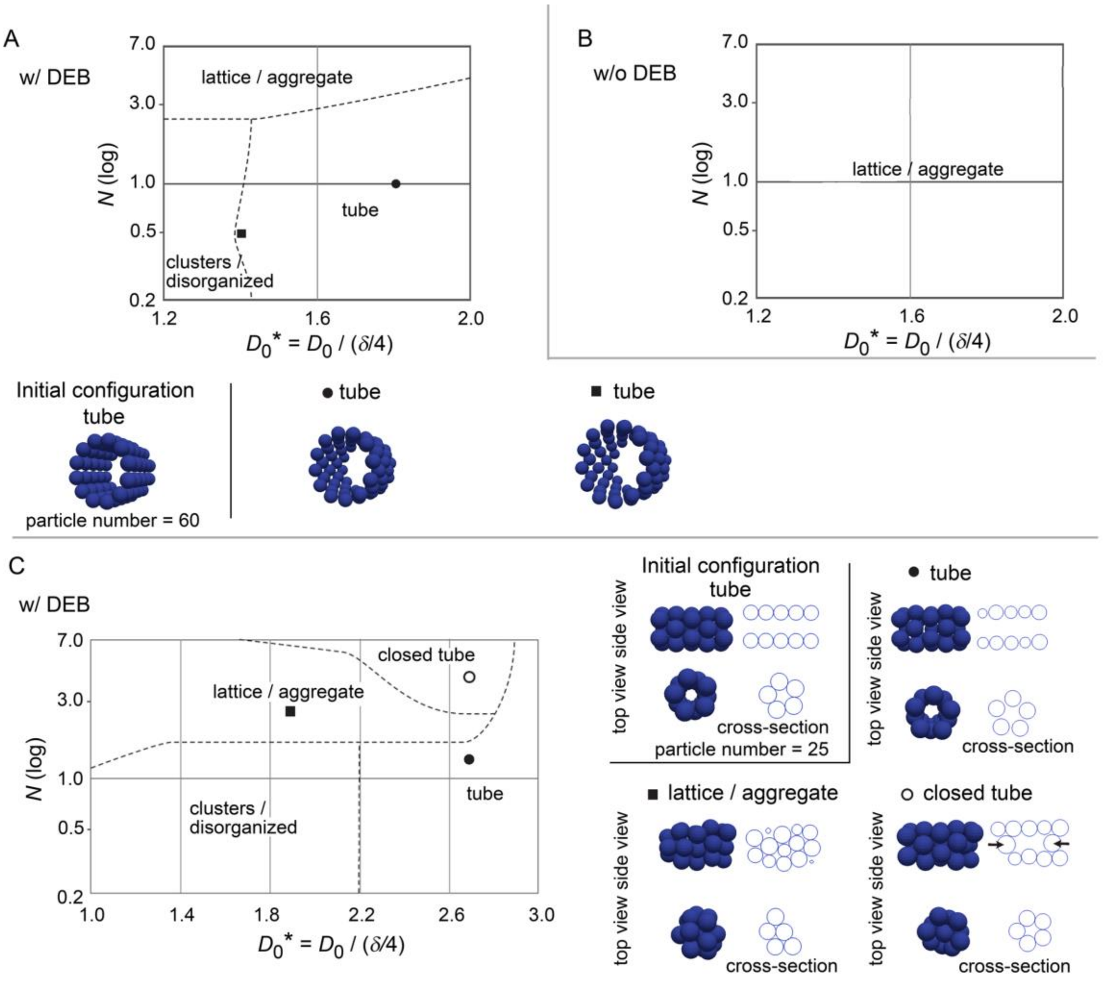


Figure S6 (related to Figure 5): Simulation outcomes and their diagrams, based on distance–force curves; starting from tube

A-B. Outcomes of simulations starting from a tube with a phase diagram are shown in a similar manner to Fig. 5. Particle number = 60.

C. Simulation outcomes similar to A except that a smaller number of particles was introduced: number of particles = 25.

The diameters of the blue spheres are equivalent to the mean diameter.

The location of the “tube” phase in the diagrams in A and C almost corresponds to that of the “cavity” in Fig. 5.

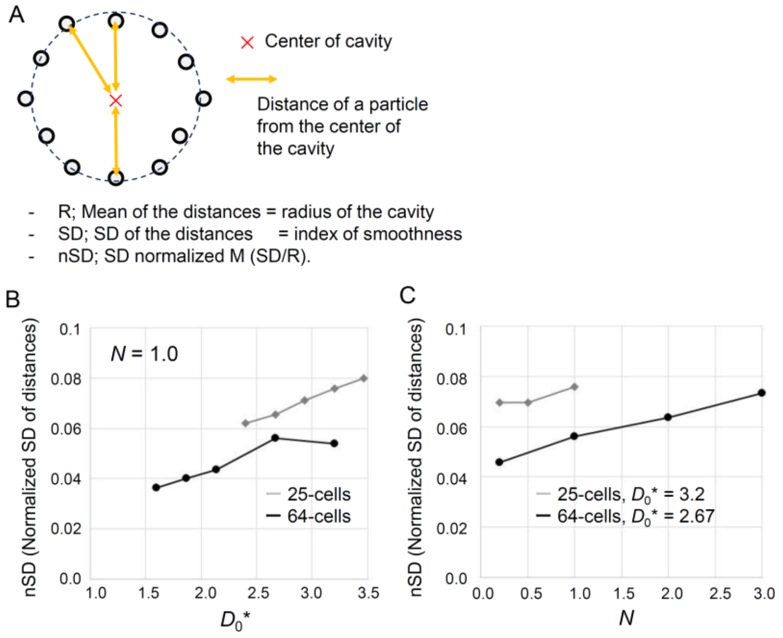


Figure S7 (related to Figure 5): Smoothness of cavities generated by simulations

A. Definition of the smoothness where a cross-section of a cavity is shown. The center of a cavity was defined as the centroid of all particles. The radius of the cavity was defined as the mean of the distances of the particles from the center of the cavity. The SD of the distances was normalized by the radius of the cavity, resulting in nSD as an index of the smoothness of the cavity. A smaller value of nSD means smoother surface of a cavity.

B-C. The smoothness (nSD) of cavities generated by simulations were evaluated. The simulation data are derived from Fig. 5B for “25-cells” and from Fig. S5A for “64-cells”. In B, $N$ was fixed at 1.0, and in C, $D_0$* was fixed at 3.2 or 2.67 for 25 or 64-cells, respectively. nSD becames smaller under the conditions of smaller values of $D_0$* and/or $N$.

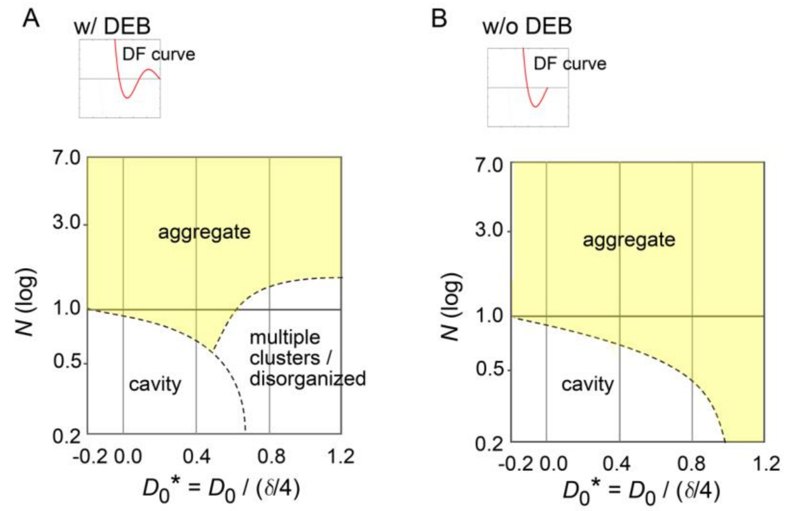


Figure S8 (related to Figure 5): Phase diagram for conditions of smaller value of $D_0$*

Phase diagrams were generated in a manner similar to Fig. 5B-C under the condition of $D_0$* < 1.2. A and B correspond to Fig. 5B and 5C, respectively. The "cavity" class equivalent to that in Fig. 6 was generated.

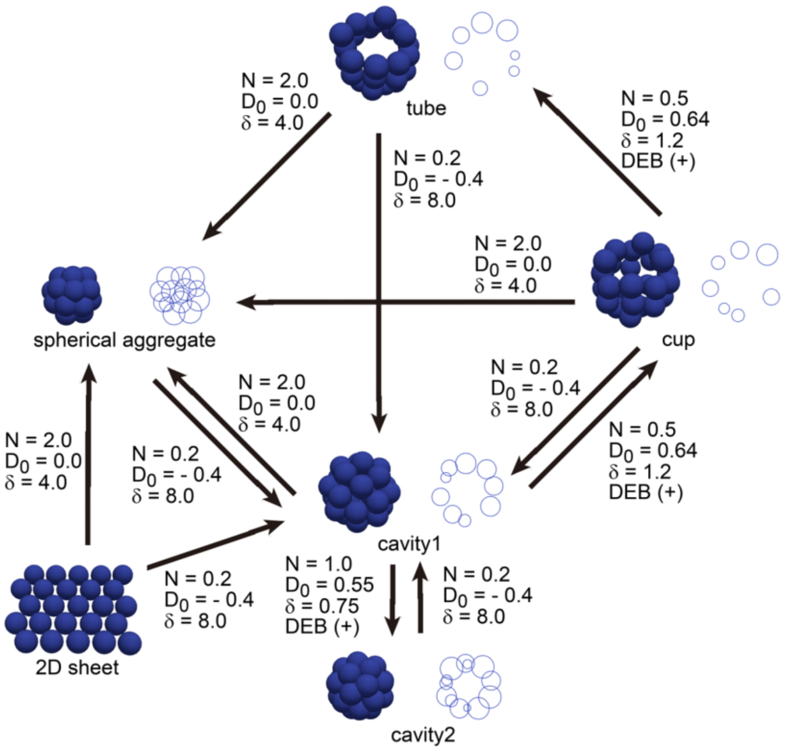


Figure S9 (related to Figure 6): Possible morphological transformation by changing distance–force curves

Under the condition of the equation 1, examples of possible morphological transformation are shown. In these simulations, the values of the parameters $N$, $D_0$, and $\delta$ were changed, and the existence of DEB was also set. These conditions are shown in the figure. The diameters of the blue spheres are equivalent to the mean diameter.

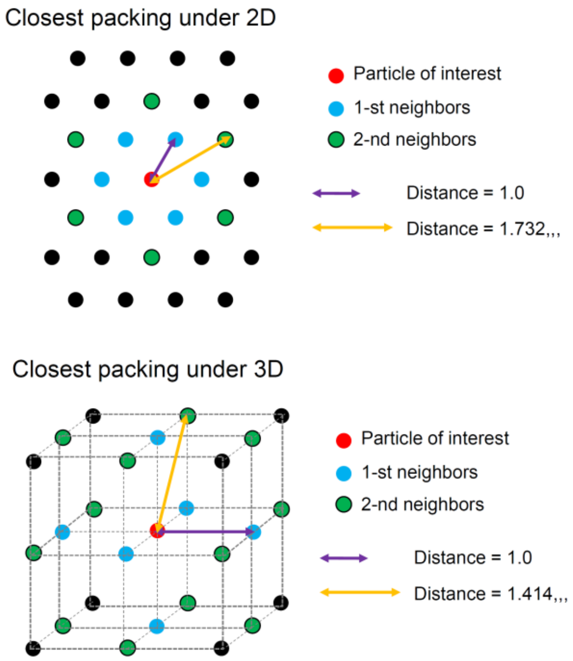


Figure S10 (related to Figure 7): Analytical evaluation of particle distributions under 2- and 3-dimensional conditions.

The closest packing situations under 2- and 3-dimensional conditions are illustrated. For the particle of interest (red circles), its $1^{st}$ and $2^{nd}$ neighbors are shown as light blue or green circles. As the relative values to the distances of the $1^{st}$ neighbors from the particles of interest, the distances of the $2^{nd}$ neighbors from the particles of interest are $3^{0.5}$ (1.732,,,) or $2^{0.5}$ (1.414,,,) under 2D or 3D conditions, respectively.

#### 2.2. Legend of Videos

Figure 2-video 1 (Movie S1): Simulations of expanding cavity.

Figure 6-video 2 (Movie S2): Stabilities of a 2D sheet and cavity-harboring structure
Whether a two-dimensional (2D) sheet and cavity-harboring structure are stably maintained under positional fluctuations was examined. 2D sheets (upper panels) correspond to Figure 6A (filled square), and cavities (bottom panels) correspond to Figure 5 (filled circle). These structures were simulated under the same distance–force (DF) curves as those used in Figure 6 and Figure 5, respectively. Positional fluctuations were given as uniform random numbers along each x-, y-, and z-axis, for each particle. The maximum displacement by the fluctuations was set to be 10% or 100% of the expected displacement provided by the maximum attractive forces of the DF curves, for each iteration during the simulations.

Figure 6-video3 (Movie S3): Simulations of possible morphological transitions shown in Figure S9 are shown.

#### 2.3. Source Data

Nuclear tracking data
Uploaded on figshare (https://figshare.com); https://doi.org/10.6084/m9.figshare.24125757

Figure 3-Source Data 1 contains following files.
Nuclear tracking data; mouse blastocyst stage (sample1-4.csv)

Figure S4-Source Data 1 contains following files.
Nuclear tracking data; MDCK cyst (sample1-4.csv)

3D force data
Uploaded on figshare (https://figshare.com); https://doi.org/10.6084/m9.figshare.24125757

Figure 3-Source Data 2 contains following files.
3D force data of cell–cell interaction; mouse blastocyst stage (4 samples: out_force_data.dat, out_xyz_data.dat)

Figure S4-Source Data 2 contains following files.
3D force data of cell–cell interaction; MDCK cyst (4 samples: out_force_data.dat, out_xyz_data.dat)

3D force map can be visualized by reading following files in the ParaView software (https://www.paraview.org/). These files will be uploaded on Web sites.
out_display_for_ParaView_line_ref[N].vtk
out_display_for_ParaView_xyz_sphere_polygon_ref[N].vtk

Data of distance–force/potential profiles
Uploaded on figshare (https://figshare.com); https://doi.org/10.6084/m9.figshare.24125757

Figure 4-Source Data 1 contains following files.
Distance–force/potential profiles; mouse blastocyst stage (DF_DP_mouse_blastocyst.xlsx)
Distance–force/potential profiles; MDCK cysts (DF_DP_MDCK_cyst.xlsx)
